# In vitro experiments and kinetic models of pollen hydration show that MSL8 is not a simple tension-gated osmoregulator

**DOI:** 10.1101/2021.10.19.464977

**Authors:** Kari Miller, Wanda Strychalski, Masoud Nickaeen, Anders Carlsson, Elizabeth S. Haswell

## Abstract

Pollen, a neighbor-less cell that contains the male gametes, undergoes multiple mechanical challenges during plant sexual reproduction, including desiccation and rehydration. It was previously showed that the pollen-specific mechanosensitive ion channel MscS-Like (MSL)8 is essential for pollen survival during hydration and proposed that it functions as a tension-gated osmoregulator. Here we test this hypothesis with a combination of mathematical modeling and laboratory experiments. Time-lapse imaging revealed that wild-type pollen grains swell and then stabilize in volume rapidly during hydration. *msl8* mutant pollen grains, however, continue to expand and eventually burst. We found that a mathematical model wherein MSL8 acts as a simple tension-gated osmoregulator does not replicate this behavior. A better fit was obtained from variations of the model wherein MSL8 inactivation is independent of its membrane tension gating threshold or MSL8 strengthens the cell wall without osmotic regulation. Experimental and computational testing of several perturbations, including hydration in an osmolyte-rich solution, hyper-desiccation of the grains, and MSL8-YFP overexpression, indicated that the Cell Wall Strengthening Model best simulated experimental responses. Finally, expression of a non-conducting MSL8 variant did not complement the *msl8* overexpansion phenotype. These data indicate that, contrary to our hypothesis and to known MS ion channel function in single-cell systems, MSL8 does not act as a simple membrane tension-gated osmoregulator. Instead, they support a model wherein ion flux through MSL8 is required to alter pollen cell wall properties. These results demonstrate the utility of pollen as a cellular-scale model system and illustrate how mathematical models can correct intuitive hypotheses.

## INTRODUCTION

Plant cells are unique mechanical systems. They have a strong, yet flexible, outer wall containing a soft, but turgid protoplast resulting in a highly pressurized cell that is often cemented to its neighbors. Turgor pressure must be controlled during growth, development, and osmotic changes in order to maintain cell and tissue integrity. Mechanical models of plant tissue development have been highly informative^1–4^. However, at the cellular scale, plant mechanobiology is affected by physical connections to neighboring cells. Wall-to-wall adhesion adds external forces and responses that complicate mechanical characterization of any one cell^5,6^. Here, we employ pollen grains as a neighbor-less model system for biomechanical characterization of plant cells.

Pollen grains undergo drastic mechanical changes throughout their normal development. Pollen, which is the male gametophyte, develops in the anthers. After meiosis and several rounds of mitosis, each grain desiccates. Some species, including the model flowering plant *Arabidopsis thaliana*, eventually contain less than 30% water^7^. Dry grains are then released from the anther to travel on pollinators or by air to reach the stigma, the receptive part of the flower. Once a compatible association is formed, the pollen grains rehydrate over the course of about 10 minutes, using moisture from the female tissue^8^. The now metabolically active pollen extends a tube structure that passes through stigmatic tissue and elongates to reach an ovule. Finally, the end of the tube bursts and releases the sperm cells in the proper location for fertilization. The entire process of sperm cell delivery to the egg requires careful control of pollen cell mechanics to prevent premature cell lysing while also maintaining rapid growth^9^. A deeper understanding of the mechanics of this process is relevant to agriculture and ecology, as all angiosperms require pollen to produce the next generation of plants. It is particularly crucial in the face of climate change, as pollen are exceptionally sensitive to high temperature^10^.

Multiple mathematical descriptions of pollen tube tip growth^11–16^ and pollen grain desiccation^17,18^ have been reported, but not pollen hydration. A few mechanisms are known to control pollen hydration, as described below, but this process has not been addressed mathematically. Important to the hydration process is the pollen grain cell wall, which is covered by a tough and water-insoluble outer layer called the exine. The exine is itself covered in a lipid and protein-based coat called the pollen coat and components of the coat are essential for establishing a productive interaction with the stigma prior to hydration^19–22^. There many other genes, such as those related to the SnRK1 complex^23,24^, that have been identified as playing a role in pollen hydration on the stigma^25^. Another feature of the pollen cell wall are apertures, areas of the wall lacking the exine layer, that allow for shrinking and expansion of the wall via folding and unfolding and can provide a route for pollen tube emergence^17,18,26,27^. Pectin in the cell wall may also contribute to hydration dynamics^28–30^. In the protoplast, aquaporins are important for water transport during in vivo hydration^31^ and the mechanosensitive (MS) ion channel MscS-Like (MSL) 8 plays a key role in maintaining pollen grain integrity during hydration and germination^32^.

MS channels are known to contribute to cell survival and/or volume regulation during hypoosmotic shock in all kingdoms of life^33–35^. This function has been explored in *E. coli* where MS channels of Small (MscS) and Large (MscL) conductance open in response to elevated membrane tension^36–38^. Upon opening, MS channels are thought to release osmolytes which will slow the entry of water, reduce cellular volume, and allow the cell to recover from osmotic shock^39,40,55^. Kinetic modeling of hypoosmotic shock in *E. coli* requires MscS and MscL channel activity to accurately simulate the observed volume changes^41^.

MSL8 may perform a similar function in pollen. MSL8 localizes to the plasma membrane and pollen lacking MSL8 shows dramatically decreased viability compared to wild type after two hours of in vitro hydration in water^32^. Moreover, when pore-blocking point mutations are introduced, MSL8 is no longer able to maintain viability during in vitro hydration^42^. These observations support the idea that MSL8 is important for pollen grain osmoregulation (i.e. acting as an “osmotic safety valve” as previously proposed^43^). However, direct evidence of ion flux through MSL8 during the early stages of hydration is lacking. Given that a close homolog of MSL8, MSL10, has a non-conducting function^44,45^, it is possible that MSL8 contributes to cellular integrity during hydration through signaling rather than through ion flux, or that ion flux through MSL8 has functions other than osmoregulation. Here, we combined experiments and mathematical modeling to test the osmotic safety valve hypothesis for MSL8 function in pollen hydration.

## RESULTS

### In vitro hydrating pollen grains lacking MSL8 continue to expand for minutes while wild-type grains stabilize within 30 seconds

To better understand the mechanics of pollen hydration and test our hypothesis that MSL8 is a tension-gated osmoregulator, we developed an assay to quantify pollen size changes during the initial stages of hydration. Fresh pollen was placed onto a glass bottom dish for imaging and a recording sequence was started directly before a drop of deionized (DI) water was applied. Most pollen grains stuck to the bottom of the dish, allowing for consistent imaging **(Figure 1A)**. By the end of hydration, WT pollen expanded ∼6.5 µm in width but shrank ∼1.5 µm in length **(Figure 1B)**. Any pollen that visibly burst was excluded from analysis.

**Figure 1.**
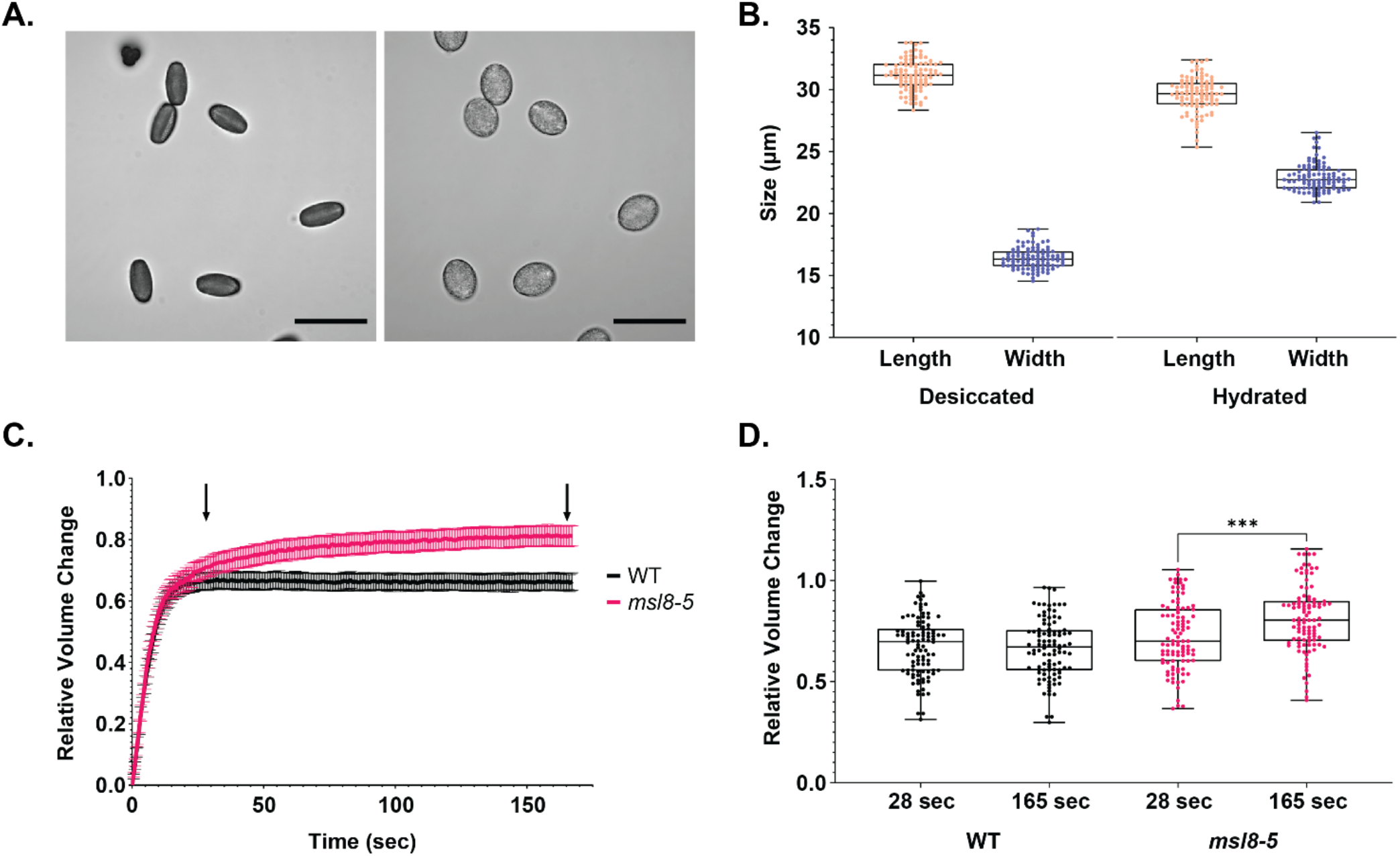
In vitro hydrating *msl8* mutant pollen grains continue to expand while wild-type grains stabilize within 28 seconds. (A) Image of Arabidopsis thaliana pollen before and after addition of DI water. Scale bars are 50 microns. (B) Length and width both before and after in vitro hydration (N=100 grains per treatment). Boxes are 1st quartile, median, and 3rd quartile while whiskers are minimum and maximum values. (C) Relative pollen grain volume over time (N=100 grains per genotype). Volume calculated assuming an ellipsoid shape and each pollen grain is normalized to itself. Bars are 95% confidence intervals (CI). (D) Comparison of the relative volume after the initial rapid hydration (the 28 second time point) and at the end of the assay (165 seconds). These time points are indicated with arrows in C. The Mann-Whitney test was used to compare *msl8-5* with itself (p<0.005) and WT with itself (p=0.70).

We took measurements every ∼0.5 seconds over the course of 165 seconds and used these dimensions to estimate the volume of an approximated 3-dimensional ellipsoid shape lying on the substrate. We then calculated the relative volume change: (initial volume – current volume)/initial volume. This value allowed us to normalize against variation in the initial desiccated grain size. WT pollen grains rapidly expanded, and after about 28 seconds of exposure to DI water, stabilized with a final volume increase of ∼60% that did not significantly change by 165 seconds (**Figure 1C-D; Supplemental Figure 1A**). We repeated this assay with pollen from an *msl8* null mutant, *msl8-5*, which was previously created via CRISPR/Cas9 gene editing^46^. We observed that *msl8* mutant lines had a high number of pollen bursting events: 12% burst compared to 1.5% in the WT (**Supplemental Figure 1B**). This was expected from previously published data showing reduced hydration survival^32,46^. While those *msl8-5* pollen grains that remained intact for the duration of the assay initially swelled with the same kinetics as WT pollen, they continued to expand throughout the time course, achieving an extra 12% expansion after initial rapid hydration. We observed this same phenotype in another *msl8* mutant line (*msl8-8*) and two *msl7-1 msl8* mutant lines (*msl7-1 msl8-6* and *msl7-1 msl8-7*)^46^ **(Supplemental Figure 1C)**. MSL7 is closely related to MSL8, but is expressed only in pollen tubes and stigma cells and is no known phenotype of *msl7-1* mutants^46^. The overexpansion during hydration observed here in *msl8* mutant pollen suggested that MSL8 is required to control the build-up of turgor in response to hypo-osmotic swelling, which follows our hypothesis that MSL8 acts as an osmotic safety valve.

### A simple kinetic model of pollen hypoosmotic swelling fails to reproduce experimental observations

To simulate the experimental data and test our assumptions about the system, we developed an ordinary differential equation that describes pollen grain expansion. This model incorporated several key properties of pollen, including an osmolyte-rich protoplast, a resilient cell wall, and MSL8 channel function. We approximated the pollen grain as a sphere and modeled the outside of the pollen grain as a single unit, with cell wall stiffness resisting internal turgor pressure. The membrane stiffness was used for calculating membrane tension, but its contribution to cell stiffness was considered negligible. Most parameter values were estimated using existing measurements in the literature **(Table 1)**^38,47–49^, but those that were unavailable were fitted to the data (see Methods).

**Table 1.**
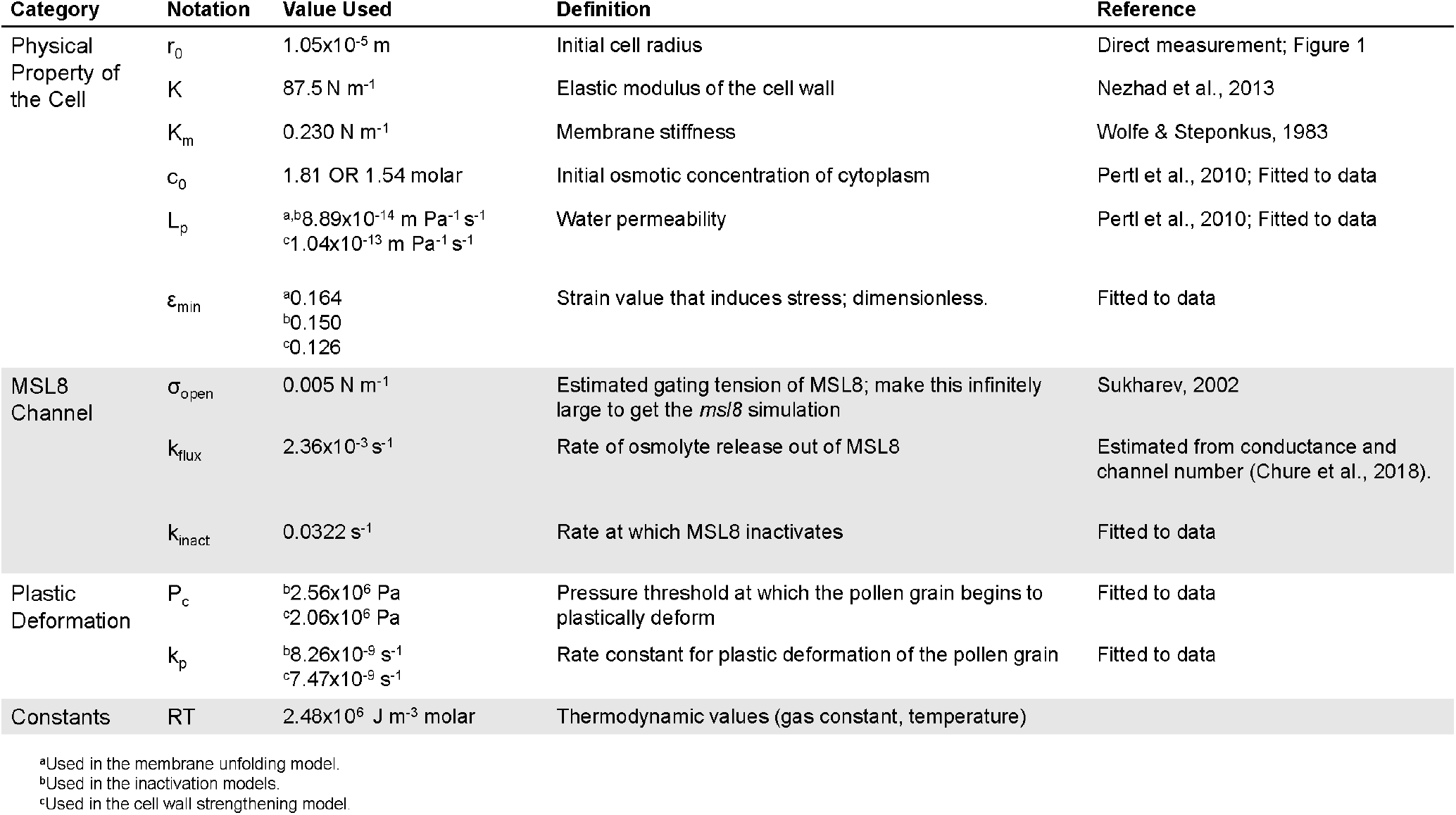
Parameters used in pollen hydration models.

Each time step of the simulation began by calculating the membrane tension (σ) based on the stiffness of the membrane and the size of the pollen grain at that point in time:

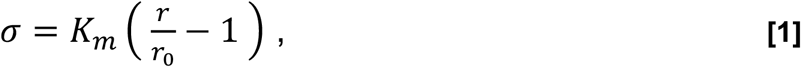

where K_m_ is the membrane stiffness, r is the radius, and r_0_ is the initial radius. Our initial model assumed that when membrane tension reaches the opening tension (σ_open_) threshold of MSL8, the channel will begin to release osmolytes. When the membrane tension drops back below the threshold, osmolyte release stops. The rate of change in osmolyte concentration is

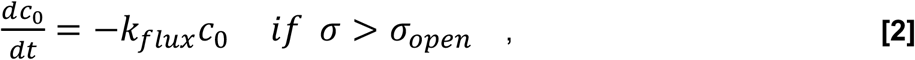

where c_0_ is the initial osmotic concentration and k_flux_ is the rate of ion flux through MSL8. Finally, we calculated the current osmolyte concentration (c) and determined the change in radius (r) as water follows the osmotic gradient and enters the pollen grain:

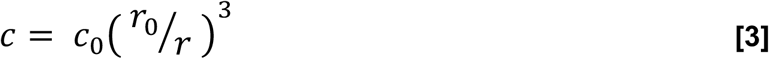

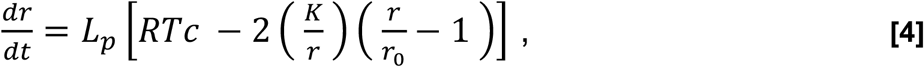

where L_p_ is the water permeability, K is the stiffness of the cell wall, R is the Boltzmann constant, and T is temperature. The second term inside the brackets represents the Laplace pressure corresponding to the membrane tension.

We ran two initial simulations: one that included MSL8 channel function (representing WT pollen) and one that removed channel function such that osmolytes never exit the grain (representing *msl8* pollen). This simulation assumed that when the gating tension is reached, all channels open to release osmolytes. We did test the idea that channels open gradually by incorporating a slow increase in the k_flux_ value after σ_open_ is reached, but it did not make any appreciable difference in the results. Thus, we retained the assumption that all channels open at once when σ_open_ is surpassed. We noticed three main discrepancies between the simulations and experimental data or the accepted values from the literature. First, the predicted membrane tension was about four times higher than the estimated membrane lytic tension of a protoplast (10 mN/m)^50^, as shown on the right Y-axis in **Figure 2A**. Second, the WT simulation showed volume increasing rapidly, peaking, and then steadily decreasing, while experimental data showed WT pollen volume stabilizing after about 30 sec. Third, the *msl8* simulation predicted a relatively rapid stabilization of the volume, but experimental data showed continued expansion for at least 150 seconds.

**Figure 2.**
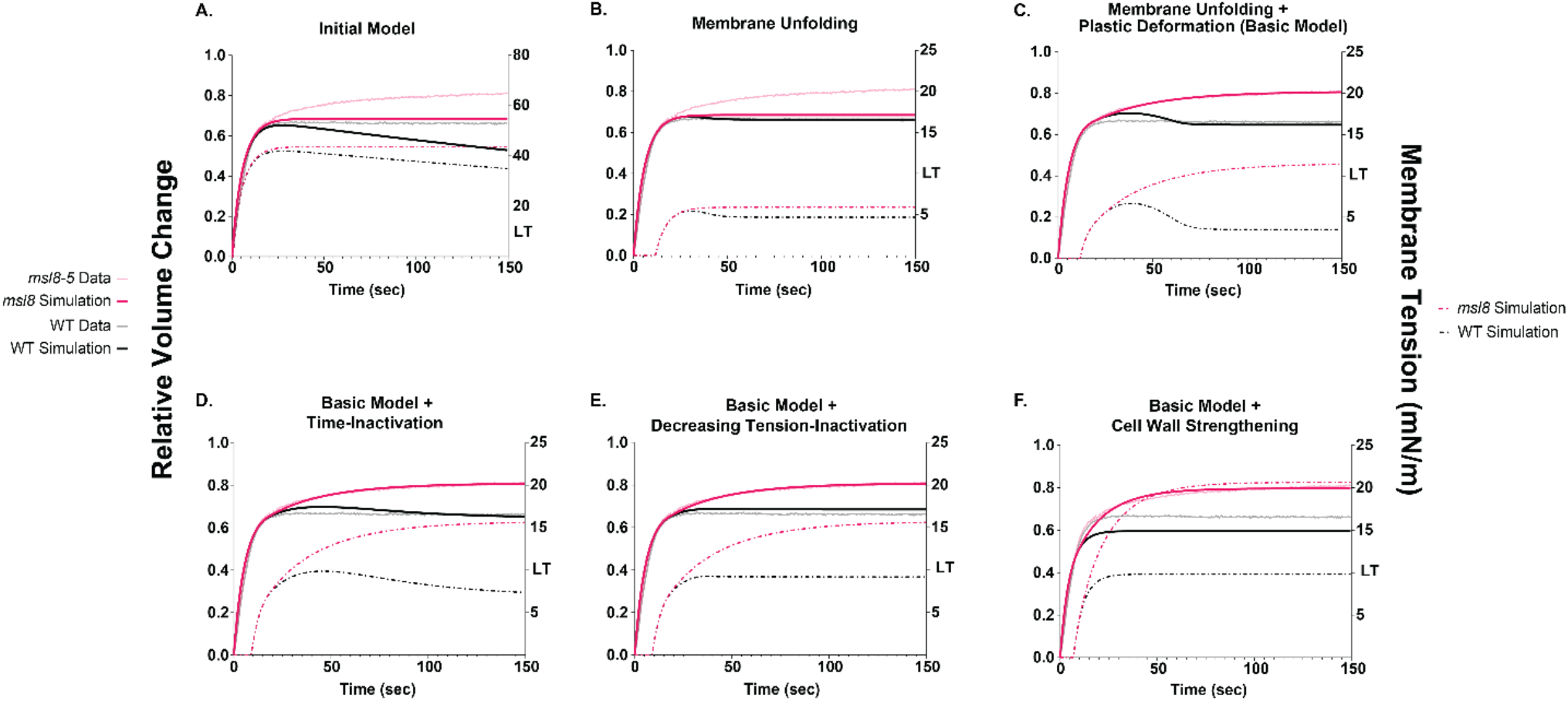
Multiple kinetic models of pollen hydration volume compared to experimental observations. (A) Simulations from the initial model for pollen hydration, which assumes that osmolyte release through the MSL8 channel begins when the tension threshold (σ_open_) is met, then channel function ceases when the tension decreases past the threshold. There is no osmolyte release in the *msl8* simulation. (B) Simulations from the membrane unfolding model, which is the initial model plus the assumption that membrane tension does not increase until the cell reaches a certain amount of strain (ε_min_). (C) Simulations from the membrane unfolding + plastic deformation model (referred to hereafter as the “Basic Model”). Plastic deformation of the cell wall when the pressure exceeds a critical threshold (p_c_). (D) The Time-Inactivation Model assumes that ion flux through the MSL8 channel function begins when the tension threshold (σ_open_) is met, then channel function ceases after a period of time (k_inact_). Plastic deformation of the cell wall is possible in both WT and *msl8* simulations. (E) The Decreasing Tension-Inactivation Model assumes that ion flux through the MSL8 channel begins when the tension threshold (σ_open_) is met, then ceases as soon as membrane tension begins to decrease. Plastic deformation of the cell wall is possible in both WT and *msl8* simulations. (F) The cell wall strengthening model which assumes MSL8-dependent cell wall strengthening, but no osmoregulation. (A-F) The estimated lytic membrane tension threshold is indicated by “LT” (10 mN/m).

### Variations on the initial model that include membrane unfolding and plastic deformation of the cell wall better reproduce experimental observations

To address the first two discrepancies, we considered possible differences between the pollen grain and other systems that show experimental volume overshoots (*E. coli*^41^ and yeast^51,52^). Freeze-fracture imaging of dry pollen^53^ suggest that the desiccated pollen membrane may not be taut, but rather has folds of extra material that allow it to expand to some extent without added tension. Simulating this effect involved calculating a strain (ε) value:

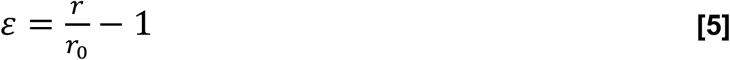

Once the strain reached a threshold (ε_min_), it was assumed in the model that all membrane reserves had fully unfolded, and membrane tension started to build:

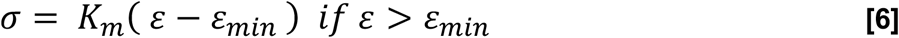

Incorporation of membrane unfolding lowered the final simulated membrane tension to physically reasonable values and removed some of the overshoot seen in the WT simulation **(Figure 2B)**. In all models hereafter, we adjusted ε_min_ to keep WT tension below the lytic level.

Nevertheless, the *msl8* simulation in the membrane unfolding model still showed a stabilized volume rather than the characteristic continued expansion seen in wet lab experiments (**Figure 2B**). Plant cell walls are sometimes described as “elastoplastic”, meaning that the wall will behave elastically until a stress threshold is reached and then permanent (i.e. plastic) deformation will occur^54^. To see if this type of material behavior could better simulate the experimental data, we incorporated plastic deformation into the model by calculating the cell pressure (p):

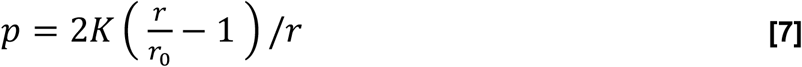

and setting a critical pressure threshold (p_c_). Only after the pressure exceeded this critical pressure would the cell wall undergo plastic deformation, determined by a deformation rate constant (k_p_):

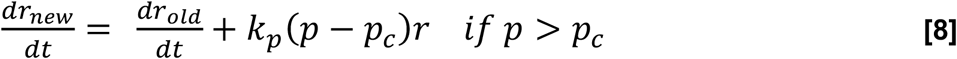

We termed this model, which incorporates both membrane unfolding and plastic deformation of the cell wall, the “Basic Model”. The Basic Model predictions aligned well with experimental observations in that the WT stabilized while the *msl8* pollen continued to expand **(Figure 2C)**. Membrane tension in the *msl8* simulation did rise above the lytic tension, but we did not consider this to be unrealistic because some *msl8* pollen did explode during the hydration process (**Supplemental Figure 1B**).

### Adding channel inactivation or MSL8-dependent effects on cell wall stiffness produce models that fit experimental data

We next tested the effect of several variations on MSL8 channel behavior on the ability of the model to produce a stable volume in WT pollen grains after ∼30 seconds of hydration. Adjusting the threshold gating tension (σ_open_) of the MSL8 channel did not produce an improved fit **(Supplemental Figure 2)**, but three other variations did. The first variation, which we call the “Time-Inactivation Model”, assumed that channel inactivation occurs spontaneously with a fixed rate. This phenomenon has been suspected to occur with other MS channels, and is a well-documented behavior of MscS^55–57^. We introduced a closing rate (k_inact_) for MSL8 that was determined by fitting to the experimental data **(Table 1)**. The closing rate was used to modify the rate at which osmolytes were released:

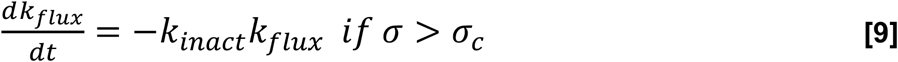

Compared to the Basic Model, the Time-Inactivation Model had improved fit to the experimental data, but it produced a small, temporary volume overshoot in the WT **(Figure 2D)**.

A second model variation, the “Decreasing Tension-Inactivation Model”, assumed that inactivation occurred as soon as the membrane tension started to decrease (but not necessarily dropping below the closing tension of the channel). MSL10 shows a related behavior in that its threshold tension and open probability are dependent on the rate at which tension is applied to the membrane^58,59^. Incorporating this modification into the Basic Model only required that k_flux_ be set to zero when the change in membrane tension 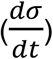 becomes less than zero. The Decreasing Tension-Inactivation Model variation fit the data well (**Figure 2E**). The curve shapes for both WT and *msl8* pollen simulations were indistinguishable from the experimental data. However, as in the Time-Inactivation Model, membrane tension in the *msl8* mutant simulation was 1.3-2 times over the estimated lytic tension of a protoplast membrane. While *msl8* mutant pollen does occasionally burst (**Supplementary Figure 1B**), most grains remain intact, suggesting that the cell wall helps support the membrane.

We next asked if MSL8 might contribute to pollen survival not directly through the release of osmolytes, but indirectly through modulation of cell wall stiffness. In the “Cell Wall Strengthening Model”, we removed all osmotic regulation by MSL8 by keeping the k_flux_ value at zero. Instead, we assumed that the presence of MSL8 channels reinforces the cell wall, thus making it more resistant to plastic behavior. This is reflected in the model by making the WT cell wall resistant to plastic deformation (i.e. omitting the plastic deformation term) while the *msl8* simulation undergoes plastic deformation past the p_c_ pressure threshold (i.e. retaining the plastic deformation term). Due to the assumption that osmolytes do not leave the pollen grain in either simulation, the c_0_ value was fitted to the WT experimental data instead of *msl8* (see Methods). Compared to the osmoregulation-based models, this model resulted in poorer accuracy in that the values do not line up with the experimental results, especially around 30 seconds into hydration (**Figure 2F**). However, the overall curve shapes were similar to the experimental results in this model. Thus, the experimental behavior of WT and *msl8* pollen grains during the first 150 seconds of hydration can be reasonably well simulated with a channel that inactivates or one that serves to strengthen the cell wall.

### Hydration in an osmolyte-rich solution restores volume stabilization in *msl8* mutant pollen and in all three model variations

To further probe the role of MSL8 as an osmoregulator during pollen hydration, we next tested the robustness of each model to experimental perturbations. We measured the response of WT and *msl8* mutant pollen to hydration in an osmolyte-rich solution, which we predicted would suppress the requirement for MSL8 to stabilize pollen volume during hydration by reducing the degree of hypoosmotic shock. We previously found that hydration of *msl8-4* pollen in a PEG 3350 solution instead of water helped restore pollen viability^32^. We replicated that approach here by hydrating the pollen in 20% (w/v) PEG 3350 and recording the first 150 seconds of hydration **(Supplemental Figure 3A-B)**. As shown in **Figure 3A**, WT pollen hydration was essentially unaffected by the addition of PEG to the hydration solution. Although the rapid initial hydration of *msl8-5* pollen was unaffected, the continued expansion in DI water was suppressed in 20% PEG. Linear regression to quantify the slope between 50 and 150 seconds of hydration indicated that, while the slope of the WT volume curve was zero in both water and PEG, the volume curve of *msl8-5* pollen in water had a slope that was significantly non-zero (p < 0.001), but zero in 20% PEG **(Figure 3B)**.

**Figure 3.**
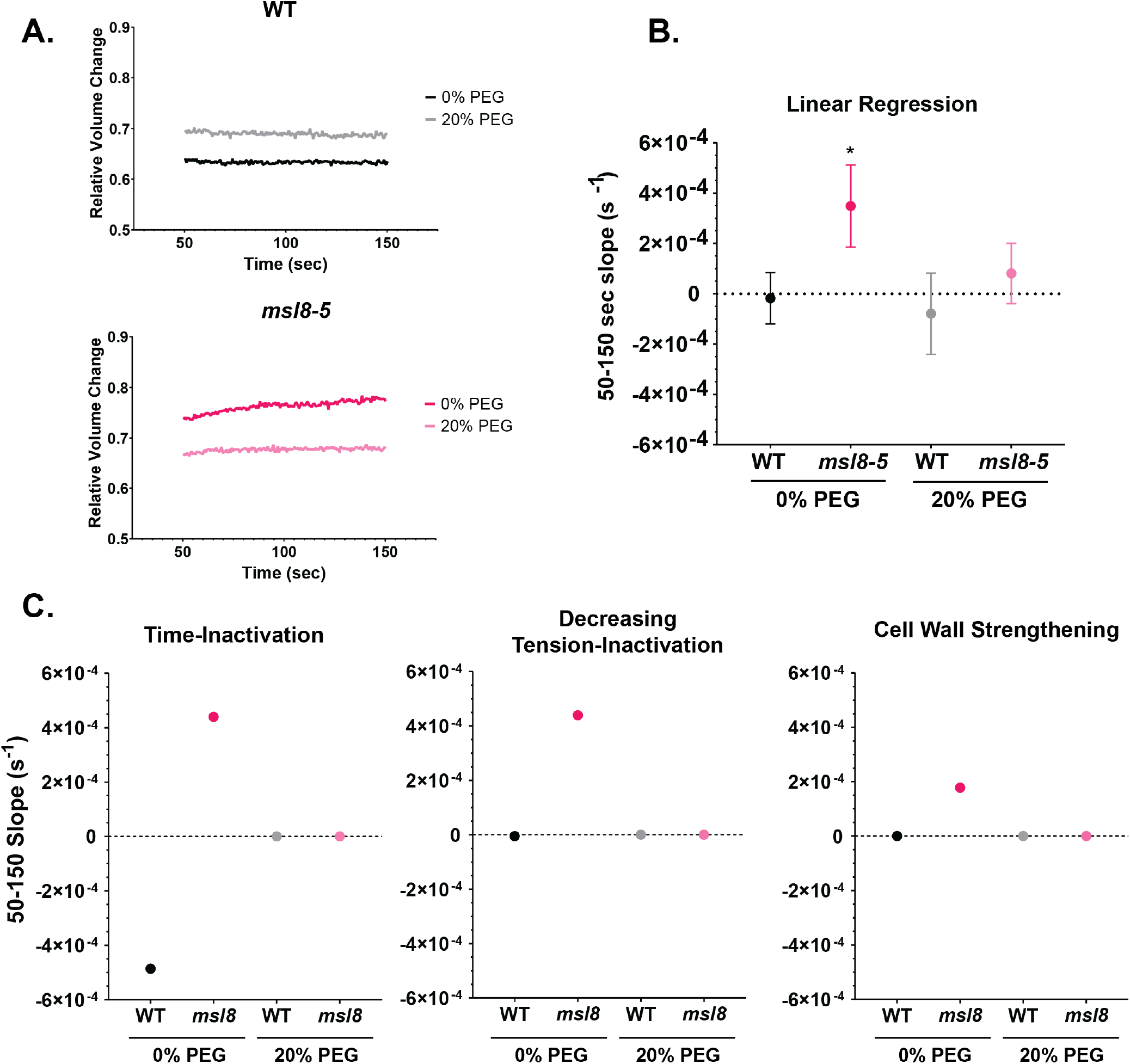
Hydration in an osmolyte-rich solution restores volume stabilization in msl8 mutant pollen. (A) The stabilization period (between 50 and 150 seconds after the addition of the hydration solution) of WT (top) and *msl8-5* (bottom) pollen grains hydrated in water (0%) or in 20% PEG 3350 (N=30 pollen grains for each genotype and treatment). Full length curves are in Supplemental Figure 3. (B) Slopes of the data in (A), estimated via linear regression for each genotype and treatment. Asterisks indicate the slope was significantly different from zero (p<0.05) which was determined via F-test. Bars are 95% CI. (C) Simulations of hydration in water and 20% PEG via modification of c_PEG_.

We next challenged our simulations with hydration in the presence of PEG. To do so, we added the parameter c_PEG_ to the change in radius calculation for the Basic Model **(eq. 4)**:

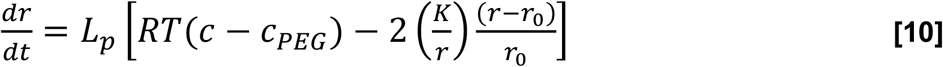

For the simulation here, the c_PEG_ value was 0.326 Osm, the predicted osmolality of a 20% PEG 3350 solution^60^. To simulate hydration in water (as in previous iterations of the model), c_PEG_ was set to 0. Simulations were carried out for the two inactivation models and the cell wall strengthening model, and the simulated slopes were calculated and compared to the experimental data **(Figure 3C, Supplemental Figure 3C-D)**. All three models predicted a stabilization of the *msl8* pollen volume in the presence of 20% PEG, consistent with our expectations and with the experimental data. In all three models, hydration in the presence of PEG resulted in a lower stabilized volume than hydration in water. However, the final pollen grain volumes in wet lab experiments were not different in the two treatments (**Supplemental Figure 3C-D**). Overall, we saw that external osmolytes could suppress the *msl8-5* expansion phenotype, and that this effect was simulated in all three variations of the kinetic model.

### The Cell Wall Strengthening model best simulated the exacerbated volume expansion of *msl8* mutant pollen resulting from additional desiccation

If, as hypothesized, MSL8 functions as an osmoregulator, increasing cytoplasmic osmolarity could exacerbate the *msl8* mutant phenotype. To test the effect of cytoplasmic osmolarity on pollen hydration, we placed freshly dehiscent pollen in a vacuum chamber overnight to more stringently dehydrate them before starting the hydration imaging assay. We found that, for the most part, this treatment had no effect on the kinetics of pollen volume changes during hydration **(Figure 4A)**. However, this extra-desiccated pollen had a slightly higher relative volume change after hydration than pollen incubated overnight at ambient conditions. For WT pollen, this slight increase was not statistically significant; however, *msl8-5* pollen swelled significantly more when extra-desiccated than when not **(Figure 4B)**.

**Figure 4.**
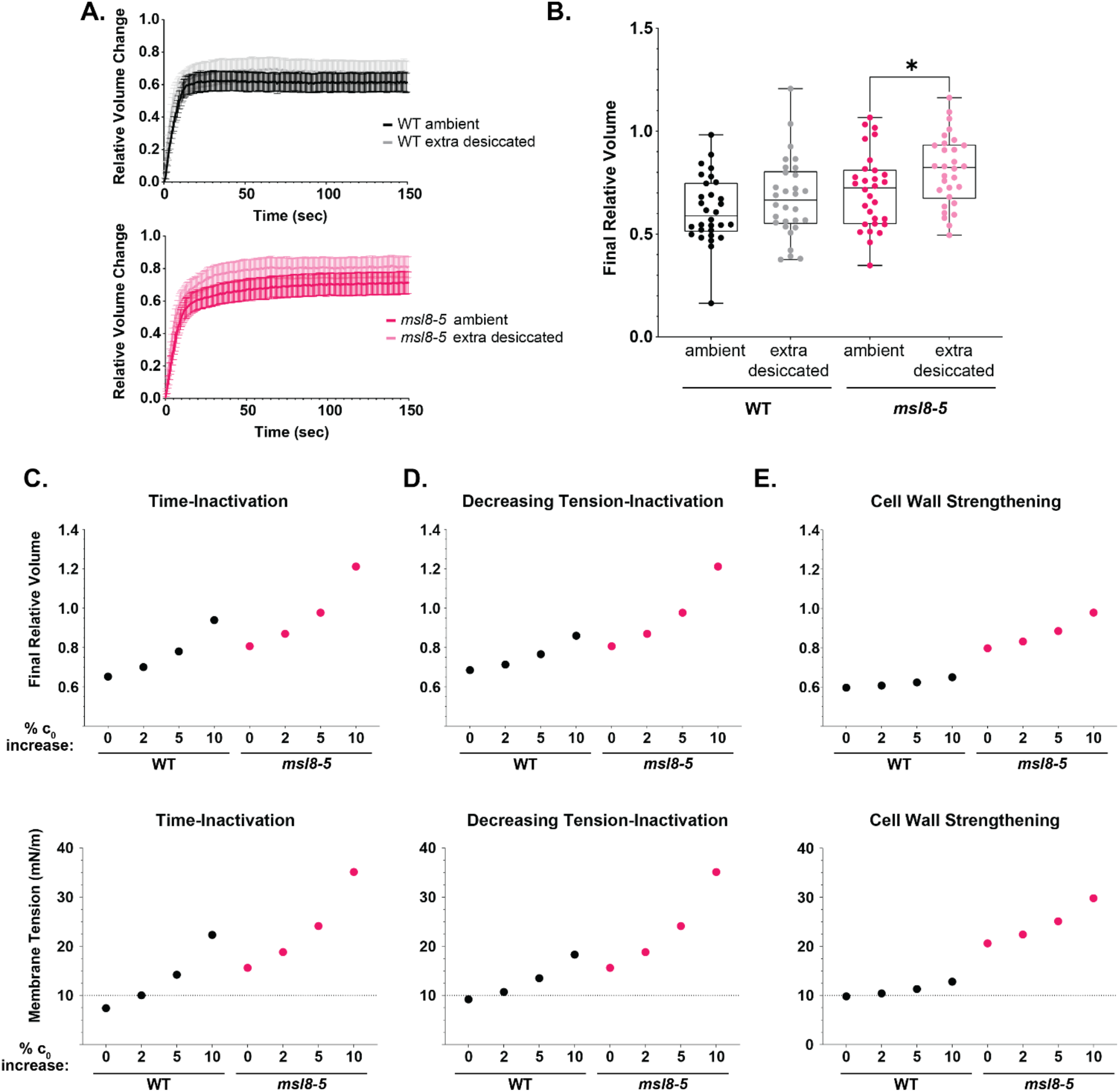
Increasing cytoplasmic osmolarity exacerbates the lack of volume stabilization in *msl8* mutant pollen during hydration. (A) Relative size change over time of hydrating WT (top) and *msl8-5* (bottom) pollen grains incubated overnight in either a vacuum chamber or ambient conditions (N = 30 pollen grains per genotype per treatment). Bars are 95% CI. (B) Final (150 s after hydration) relative volume change for pollen grains in the experiment shown in A. Mann-Whitney U test performed between the extra desiccated and ambient treatment for each genotype. Grubbs test for outliers did not identify any outliers. *msl8-5* p-value = 0.042. WT p-value = 0.22. (C-E) Final relative volume and membrane tension in simulations after altering c_0_ in the indicated kinetic models.

To simulate the effect of extra desiccation in the model variants, the osmotic concentration (c_0_) parameter was increased **(Supplemental Figure 4)**. We found that increasing c_0_ by 10% resulted in extremely high membrane tension. Thus, we tested 2%, 5% and 10% increases in c_0_ and examined the predicted final relative volume and membrane tension. In Time-Inactivation Model simulations, and to some degree the Decreasing Tension-Inactivation Model simulations, *msl8* and WT pollen increased both final volume and membrane tension with increasing c_0_ **(Figure 4C-D)**. However, Cell Wall Strengthening Model simulations showed increased swelling and tension in *msl8* pollen with increasing c_0_, while WT pollen did not change appreciably **(Figure 4E)**. Thus, increasing cytoplasmic osmolarity through extra desiccation exacerbated the lack of volume stabilization in *msl8* mutant pollen, but not in the WT, as predicted by our hypothesis that MSL8 functions as an osmoregulator during hydration. The Cell Wall Strengthening Model was the best at simulating this effect, suggesting that cell wall extensibility rather than osmolyte release could be the key difference between WT and *msl8* mutant pollen.

### Overexpressing MSL8-YFP does not affect pollen volume stabilization and this was best replicated in the Decreasing Tension-Inactivation and Cell Wall Strengthening Models

Next, we examined the effect of increased MSL8 channel number on pollen grain volume during hydration. If MSL8 functions as an osmotic safety valve, we would expect that more channels would release more osmolytes and result in a lower final volume after pollen hydration. We therefore overexpressed MSL8-YFP via the pollen-specific *LAT52* promoter (*pLAT52::MSL8-YFP*)^32,61^ in the Col-0 background. We identified four heterozygous overexpression lines (OE 13, 17, 20, 27) and confirmed transgene expression via YFP fluorescence. We were unable to identify homozygous lines, likely due to previously documented negative effects of MSL8 overexpression on male fertility^32,42^. We noted that pollen grains over-expressing MSL8-YFP were smaller than WT pollen **(Supplemental Figure 5)**. Yet, when the relative volume change of pollen over-expressing MSL8-YFP was calculated – thus, the volume was normalized – the resulting curves were indistinguishable from volume curves of WT pollen **(Figure 5A)**.

**Figure 5.**
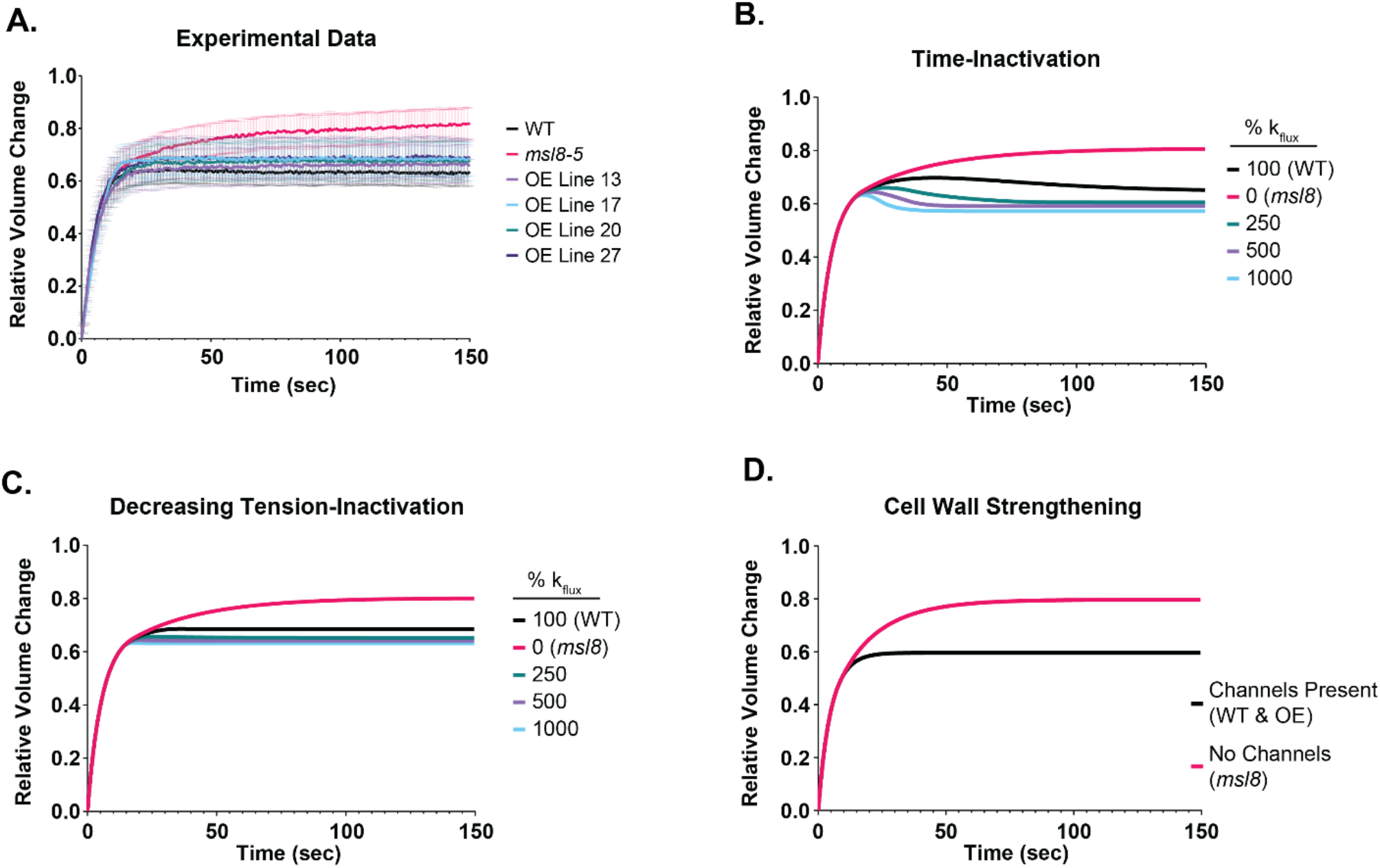
Overexpressing *MSL8-YFP* does not affect volume stabilization during pollen hydration. (A) Relative size change over time of hydrating pollen grains overexpressing (OE) *MSL8-YFP* (N=30 grains per genotype). Bars are 95% CI. (B-C) Time-Inactivation and Decreasing Tension-Inactivation model results with varying relative values of the k_flux_ parameter. (D) Cell wall strengthening model results. Note that there is no effective osmolyte release (i.e. no k_flux_ parameter), thus this result is the same as in Figure 2F.

To include MSL8 overexpression in the models, effective channel activity was increased by modifying the channel ion flux rate, k_flux_. We set 0% channel function to be equivalent to *msl8* while 100% channel function was equivalent to WT. Further increases in k_flux_ represented MSL8 overexpression. Both the Time-Inactivation Model and the Decreasing Tension-inactivation Model were relatively insensitive to k_flux_, so channel function could be increased up to 1000% without resulting in a relative volume change lower than 0.5, which is the lower end of the 95% confidence intervals **(Figure 5B-C)**. The Time-Inactivation Model showed an overshoot and recovery when channel function was high, which was not reflected in the experimental data (**Figure 5A**). However, the Decreasing Tension-Inactivation Model fit the experimental data well, with very little effect from increased channel function. The Cell Wall Strengthening Model does not have channel function so there was no k_flux_ value to increase, and therefore by definition matched this experimental result **(Figure 5D)**. To summarize, we found that, unexpectedly, overexpressing MSL8-GFP did not alter the kinetics of swelling in hydration experiments. Furthermore, two of our models (Decreasing Tension-Inactivation and Cell Wall Strengthening) simulated the experimental data well, suggesting that MSL8 does not function as a simple tension-gated osmoregulator.

### Pore-blocked MSL8 channels are unable to prevent overexpansion during pollen hydration

Thus far, our initial hypothesis that MSL8 functions as an osmoregulator was not supported so we next tested to see if ion flux through MSL8 was required for volume stabilization. To do this, we used a previously characterized MSL8 point mutation (MSL8^F720L^) that abolishes ion conductance and is required for pollen to survive hydration^42^. MSL8-GFP or MSL8^F720L^-GFP were expressed from the genomic context in the *msl8-5* background using previously described transgenes (*gMSL8-GFP* and *gMSL8*^*F720L*^*-GFP*)^42^. Three lines were selected for each transgene and stable, full-length protein expression was confirmed via immunoblot **(Supplemental Figure 6A)**. Phenotypes were assessed using the initial hydration assay **(Supplemental Figure 6B)**. As expected, a transgene harboring the genomic version of *MSL8* complemented the *msl8-5* phenotype, as *msl8-5 gMSL8-GFP* pollen stabilized in volume between 50-150 seconds of hydration (**Figure 6A**). Quantification showed a slope that was not significantly different from zero in all three lines (**Figure 6B**). Meanwhile, *msl8-5 gMSL8*^*F720L*^*-GFP* pollen continued to expand, similar to *msl8-5* pollen, and showed a non-zero slope between 50-150 seconds (**Figure 6A-B**). Thus, osmolyte conductance through MSL8 is necessary for the volume stabilization seen in wild-type pollen.

**Figure 6.**
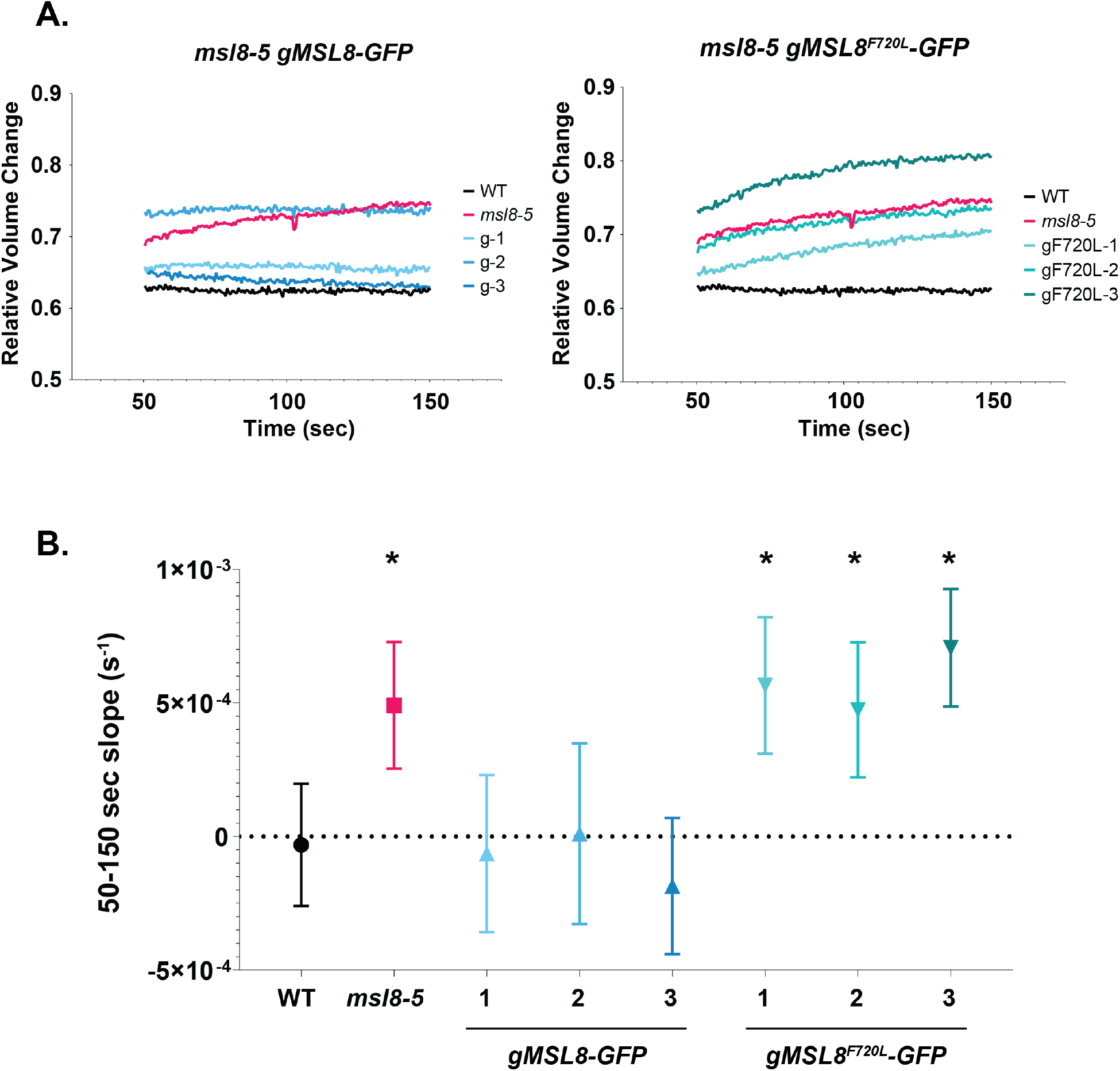
MSL8 channel conductance is required for pollen grain volume stabilization. (A) Experimental hydration results of pollen expressing *gMSL8-GFP* (left) and *gMSL8F720L-GFP* (right) (N=30 grains for each). Only the relative volume between 50 and 150 seconds is displayed as this section was used to estimate a slope; full length curves are shown in Supplementary Figure 6. (B) Slopes estimated via linear regression for each genotype. Asterisks indicate that the slope was significantly different from zero (marked with a dashed line) which was determined via F-test (p<0.05). Bars are 95% CI.

## DISCUSSION

Plant cell mechanics is complicated by the presence of a cell wall and the generation of multiple atmospheres of turgor pressure. The process of in vitro pollen grain hydration represents a relatively simple starting place for modeling plant cell mechanics–due to the absence of neighbors, to the isotropic nature of the expansion, and to the established requirement for the MS ion channel, MSL8, for survival. Here, we characterized the kinetics of pollen swelling during the first 150 seconds of pollen hydration in a range of osmotic conditions and MSL8 levels. In addition, we developed and tested several related models of pollen hydration which incorporated our assumptions about cell wall mechanics, membrane mechanics, and MS channel function. Our primary goal was to test the hypothesis that MSL8 acts as an osmotic safety valve to regulate pollen volume during the hypoosmotic shock of hydration. Taken together, our results do not support this hypothesis. While a model using our initial assumptions predicted the volume “overshoot” that is seen in other single cell systems responding to hypoosmotic shock, including *E. coli*^41^ and yeast^51,52^ **(Figure 2A)**, this was not observed in our experimental results **(Figure 1C)**. Subsequent modifications of the initial model led us to a Basic Model that 1) presumes there are membrane reserves that unfold during hydration, which impacts the timing of MSL8 channel opening **(Figure 2B)** and 2) that the cell wall behaves as an elastoplastic material **(Figure 2C)**. Because this model still produced a transient volume overshoot that is not seen in the actual data, we generated three alternative modes for MSL8 channel function, all of which removed the overshoot and successfully simulated the response of pollen to hydration in water: 1) Time-Inactivation, 2) Decreasing Tension-Inactivation, or 3) Cell Wall Strengthening **(Figure 2D-F**). To distinguish between these models, we determined the effect of osmotic support in the hydrating solution, increasing cytoplasmic osmolarity through extra desiccation prior to hydration, and channel overexpression, then compared experimental results to predictions from all three model variants. While no model produced a perfect simulation of the experimental results, the Cell Wall Strengthening Model was the best across all three perturbations.

### Does desiccated pollen have membrane reserves?

It is likely that the desiccated state of the pollen grain adds intricacies to its mechanics, and that this would affect the function of any embedded proteins, including MS ion channels. First, it has long been proposed that the lipid bilayer in desiccated pollen is not intact^62–64^. However, recent freeze-fracture electron microscopy studies indicate a clear membrane around the outside of the pollen protoplast during the earliest phases of hydration^53,65^. Second, even if they are intact, desiccated membranes may not be under tension. Dry pollen from multiple species show ruffles in freeze-fracture electron micrograph images^53,65^. Folds in the plasma membrane of a desiccated pollen grain, or during the early stages of hydration, would delay the build-up of membrane tension and therefore delay MS ion channel activity. Incorporating this phenomenon into our model kept the predicted membrane tension at a realistic value while also removing much of the volume overshoot, producing a model that was overall more accurate. We note that unexpectedly large membrane reserves have been previously documented in animal cells^66–68^ and may serve as a universal mechanism for preventing or delaying the activation of mechanosensitive processes.

### Elastoplastic behavior and swelling of the hydrating pollen cell wall

Plant cell wall material is often described as “elastoplastic”, meaning it is resistant at low forces and yielding at high forces^54^. This material property is the result of many cell wall components interacting and each contributing strength and/or flexibility^35^. A pollen hydration model that assumed the wall is a simple elastic material did not capture the volume dynamics we observed experimentally. In particular, we were unable to simulate the slow expansion seen in *msl8* pollen (compare red lines in **Figure 2A-B**). By assuming that the cell wall permanently deforms past a pressure threshold, we were able to simulate *msl8* experimental results in the Basic Model **(Figure 2C)**. While this did create a small overshoot in the WT simulation of the Basic Model, in the Time-Inactivation, Decreasing Tension Inactivation, and Cell Wall Strengthening model variations, the pressure threshold was not reached in the WT simulation; thus, there was no deformation of the cell wall and the volume remains stable, aligning with our experimental results **(Figure 2D-F)**.

Another place where the cell wall may have behaved differently from our original assumptions was when the pollen was hydrated in PEG solution. Even with plastic deformation added, our model overestimated the effect of PEG in the hydration media **(Figure 3, Supplemental Figure 3)**. Here the experimental data showed WT pollen reached the same final volume when hydrated in water as in 20% PEG, but our model predicted a much smaller final volume. The cause of this discrepancy is not clear, but it is possible that cell wall swelling or hydrogel formation could affect the observable volume. Absorbent pectin gels could be causing a wicking action that absorbs more water than our model is able to account for^28,29,69^. A swelling cell wall might also have implications on other mechanical components by reducing water entry into the protoplast which would keep membrane tension low.

### MSL8 does not function as a simple osmotic safety valve during pollen hydration

As discussed in the introduction, our initial characterization of MSL8’s role in maintaining pollen viability led us to hypothesize that the channel is acting as an osmotic safety valve to prevent overexpansion, similar to MscS in *E. coli*^32,39–42^. A pore-blocked MSL8 variant was unable to rescue the *msl8* phenotype, confirming that ion flux through the channel is necessary in some capacity **(Figure 6)**. However, our Basic model, which assumed that MSL8 is acting as an osmoregulator, did not predict the experimental result **(Figure 2C)**. We thus developed three variations of the model: 1) MSL8 inactivates after some time; 2) MSL8 inactivates when the tension starts to decrease; or 3) the key aspect of MSL8 channel function is not osmoregulation, but cell wall stiffening. Below we discuss these three possibilities in the context of existing literature on MS channel dynamics and cell wall mechanics.

#### 1) Time-Inactivation or Decreasing Tension-Inactivation of MSL8

The Initial and Basic Models assumed that osmolyte release by MSL8 channels stopped only when membrane tension dropped below the opening tension threshold of the channel, but neither replicated WT volume dynamics properly. Modeling channel inactivation after time passed (Time-Inactivation, **Figure 2D)** or after tension began to decrease (Decreasing Tension-Inactivation, **Figure 2E)** either partially or fully recapitulated the volume stabilization seen in WT pollen, respectively. Both models successfully predicted the effects of PEG hydration and extra desiccation on *msl8* pollen. The Time-Inactivation Model did not successfully predict the effect of MSL8 overexpression, and neither model predicted the lack of effect of extra desiccation on WT pollen. Furthermore, time-inactivation has not been observed for MSL8^32^, nor does it appear to close quickly as tension is released. Instead, MSL1^70^, MSL10^59^, and MSL8^32^ all close much more slowly than they open; in some cases, channels opened spontaneously after all tension was released, thereby maintaining an extended open state. Thus, channel inactivation seems unlikely explanation for our experimental data, but a more detailed study of MSL8 channel opening, closing, and inactivation kinetics will be necessary to rule them out.

#### 2) Cell wall strengthening by MSL8

The second way to achieve volume stabilization after the early stages of hydration is to strengthen the cell wall so it resists overexpansion, as shown in the Cell Wall Strengthening model **(Figure 2F)**. This simple model had the fewest parameters, successfully predicted the effect of hydration in PEG (**Figure 3C**) and extra desiccation on both WT and *msl8* pollen (**Figure 4E**), as well as the hydration phenotype of pollen overexpressing MSL8-YFP **(Figure 5)**. How might ion flux through MSL8 affect cell wall properties? MSL8 is likely to transport Cl^- 32^, so its activation would increase anions in the apoplast, depolarize the membrane, and could alter pH in the cell wall. Several components in the cell wall, such as cellulose, hemicellulose and pectin, have complex electrostatic interactions that impact the stiffness of the cell wall^71^. For example, electrostatic interactions between negatively charged pectins and positively charged extensin proteins^72^ may be affected by the ionic strength of the apoplast. Weakening the cell wall through defects in cellulose deposition did not reproduce the *msl8* mutant phenotype **(Supplemental Figure 7)**, suggesting that MSL8 could affect a non-cellulosic component of the cell wall. Future studies of the effect of MSL8 on cell wall composition (e.g. through glycome profiling) and strength (e.g. through atomic force microscopy) will be crucial to experimentally test this intriguing model.

## Conclusions and Future Directions

The data presented here reveal that MSL8 stabilizes pollen volume during the initial stages of hydration, but not via simple osmoregulation as we originally hypothesized. Rather, it suggests that *msl8* mutants have altered pollen cell wall properties, or that the MSL8 channel exhibits unusual inactivation behavior. This work highlights the utility of mathematical modeling for testing assumptions in proper physical context while also developing new, testable hypotheses. We demonstrated that our assumption of MSL8 function – which was based on MS channel function in other systems – was not entirely correct. Future computational work should incorporate the ellipsoid shape of the pollen grain, add cell wall heterogeneity like apertures, and eventually address the polarized nature of hydration and expansion that occurs in vivo and during tube germination. Overall, we believe that this model of single plant cell mechanics will be useful as we seek to understand how osmotic regulation and cell wall integrity are influencing one another.

## MATERIALS AND METHODS

### Accession Numbers

The accession numbers for the genes studied here are: *MSL8* (At2g17010), *MSL7* (At2g17000), *FRA1* (At5g47820), *CMU1* (At4g10840), and *CMU2* (At3g27960).

### Plant Materials and Growth Conditions

*Arabidopsis thaliana* plants of the Columbia-0 ecotype were used in all experiments. The *msl8-5, msl8-8, msl7-1msl8-6* and *msl7-1msl8-7* mutant lines were generated via CRISPR/Cas9 gene editing in WT or the *msl7-1* T-DNA mutant backgrounds, as described in Wang et al., 2021^46^. Seed was surface-sterilized using vapor-phase chlorine for six hours before being placed on Petri dishes containing 1/2X Murashige and Skoog (MS) medium, pH 5.7. For transgene selection, phosphinothricin (GoldBio) was added to the MS media. The plates were incubated for two days at 4°C then transferred to a 24-hour light chamber (Conviron) with 120 m^−2^ s^−1^ photons at 21°C and 50% relative humidity for 5-6 days. Seedlings were then transferred to soil and grown under 150 m^−2^ s^−1^ photons light intensity in a 16/8-hour light/dark chamber (Conviron) at 21°C.

### Pollen Hydration Imaging and Analysis

Both time lapse and fluorescence imaging were performed on an Olympus FV3000 confocal laser scanning microscope. For time lapse imaging, dry pollen was placed onto a glass bottom dish (MatTek P35G-1.5-14-C) by gently tapping 5-8 freshly opened flowers onto the glass. In some experiments, dishes were immediately used for imaging while in others, the dishes were placed into a vacuum chamber for a 12 h desiccation treatment or incubated on the benchtop for ambient treatment. Once on the microscope, pollen grains were imaged using a 20X objective. Recording began before water was added, and images were taken every 0.55 seconds over the course of hydration with either deionized water or 20% (w/v) PEG 3350. Image analysis was done using particle analysis in FIJI^73^. Only pollen grains that were not touching other grains/debris and did not visibly lyse were included in the analysis. Statistical tests were performed using GraphPad Prism Version 9.1.

### Model Fitting Procedures

All model code was run in MATLAB Version 9.4 (R2018a). To fit c_0_, the value was solved for from the steady state equation:

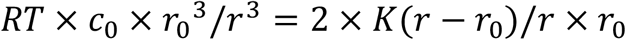

For the basic and inactivation models, the initial radius 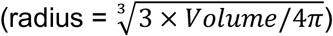 and inflection point radius taken from the *msl8-5* data was used to account for loss of osmolytes due to channel function in the WT. For the cell wall strengthening model, the initial and final radius taken from the WT data was used.

After determining the c_0_ value, the L_p_ value was fitted to the data by solving:

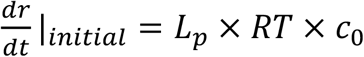

The initial dr/dt was determined from the relevant data set by averaging the change in radius every two seconds over the first twenty seconds, then taking the exponential fit (*msl8-5* initial dr/dt = 0.38; WT initial dr/dt = 0.45). For other parameter values fitted to the data (see Table 1), goodnessOfFit() with Mean Squared Error cost function was used for iterative searching of the minimized error.

### Constructs and Plant Transformation

To create pollen-specific MSL8-YFP overexpression lines, *pLAT52: MSL8-YFP*^32^ was introduced into Col-0 plants using *Agrobacterium tumefaciens* GV3101 strain-mediated transformation via floral dip. Confirmation of expression was determined through fluorescence imaging of pollen grains (488 nm excitation, 500-550 nm range detected).

To create lines expressing MSL8 and MSL8^F720*L*^ at endogenous levels, *pMSL8: MSL8-GFP* or *pMSL8: MSL8*^*F720L*^*-GFP*^42^ were introduced into *msl8-5* plants using *Agrobacterium tumefaciens* GV3101 strain-mediated transformation via floral dip. Due to low expression from the *MSL8* native promoter, immunoblotting was used to confirm the presence of full-length MSL8-GFP protein. For plants expressing *GFP* under the *MSL8* promoter, we isolated pollen via centrifugation of ∼60-80 flowers in 1 mL of water. The flowers and water were removed before exposing to two freeze-thaw cycles in liquid nitrogen. The pollen was then resuspended in 90 µL of 2X sample buffer (0.1 M Tris-HCl, 4% SDS, 20% Glycerol, 0.2% bromophenol blue, and 2% β-mercaptoethanol). 20 µL of each sample was loaded and the proteins were resolved using 10% SDS-PAGE resolving gel followed by transfer to a polyvinylidene difluoride membrane (BioRad) at 100 mA for 12 h. After blocking in 5% milk TBS-T, membranes were incubated in anti-GFP (Takara Bio, 1:5000 dilution) antibodies followed by a 2 h incubation in secondary goat anti-mouse-HRP (1:10,000 dilution; Millipore). Detection was performed using the SuperSignal West Femto Detection Kit for anti-GFP. Afterwards, the blot was stripped and re-probed using the same protocol with anti-tubulin (Sigma, 1:20,000 dilution) antibodies. SuperSignal West Dura Detection Kit was used to detect anti-tubulin (Thermo Fisher Scientific).

## SUPPLEMENTARY DATA

**Supplemental Figure 1: In vitro pollen hydration curves of all *msl8* mutant and WT lines**.

(A) Hydration curves for two wild type lines. WT Col-0 is the original parent line and WT sibling was segregated from the *msl7-1msl8-7* line. N = 30 grains for each genotype; Bars are 95% confidence intervals (CI). (B) Percentage of pollen grains that visibly burst during the first 150 seconds of hydration (N = 206 grains for WT; N = 234 grains for *msl8-5*). (C) Hydration curves from all *msl8* mutant lines.

**Supplemental Figure 2: Effects of altering the channel opening membrane tension threshold**.

(A) Volume output of simulations testing σ_open_ values of 0, 2.5, 5, 7.5, and 10 mN/m. Note that 5 mN/m is the estimated value based on existing literature. (B) Membrane tension output from the same five simulations. The dashed line marks the estimated lytic tension of the membrane.

**Supplemental Figure 3: Hydration in an osmolyte-rich solution experimental observations and simulations**.

Relative volume change of WT (A) and *msl8-5* (B) pollen hydrated in water or 20% PEG 3350 solution (N = 30 pollen grains for each genotype/treatment). Bars are 95% CI. Simulation results for both WT (left) and msl8 (right) hydration in 0% PEG and 20% PEG (assuming the MSL8 channels inactivate after a period time (C), MSL8 channels inactivate when tension starts to decrease (D), or the presence of the MSL8 channels strengthens the cell wall (E).

**Supplemental Figure 4: Hydration simulations of pollen with increased cytoplasmic osmolarity**.

Simulation results for both WT (left) and msl8 (right) pollen grains with increasing initial osmotic concentration values assuming either the MSL8 channels inactivate after a period time (A), MSL8 channels inactivate when tension starts to decrease (B), or the presence of the MSL8 channels strengthens the cell wall (C).

**Supplemental Figure 5: Volume of pollen overexpressing MSL8-YFP**.

Initial pollen volume (left) and final volume (right) of WT, *msl8-5*, and four *MSL8-YFP* overexpression lines (OE) (N = 30 grains for each). Bars are mean with 95% CI. One-way ANOVA test with multiple comparisons was used to compare each genotype among initial volume and final volume groups. Letters indicate a significant difference (p < 0.05). Grubbs test for outliers was conducted on all genotypes. One outlier from WT final volume and two outliers from OE 20 final volume were removed. Excluding the outliers did not affect the ANOVA results.

**Supplemental Figure 6: Characterization of MSL8 lines**.

(A) Immunoblot confirmation of protein presence in *msl8-5* lines expressing genomic *MSL8-GFP* or *MSL8-GFP* with the point mutation, F720L, which eliminates channel function. Protein blot was probed with anti-GFP (top). The blot was then stripped and re-probed with anti-tubulin (bottom). (B) Complete hydration curves of the lines.

**Supplemental Figure 7: Hydration of pollen cell wall mutants**.

Hydration curves of WT and *msl8-5* controls (left) and three cell wall mutants (*fra1-5, cmu1 cmu2*, and *fra1-5 cmu1 cmu2*) (right) (N=30 grains per genotype). Bars are 95% CI. All hydrations were performed in 10% PEG due to the high rate of explosions of *fra1 cmu1 cmu2* pollen.

## ACKNOWLEDGEMENTS

We thank Ram Dixit (Washington University in St. Louis) for providing *fra1 cmu1 cmu2* mutant seeds.

This work was supported by NSF Graduate Research Fellowship (DGE-2139839 and DGE-1745038) to K.M., NSF MCB 1929355 to E.S.H and A. C, and the NSF Center for Engineering Mechanobiology NSF CMMI 1548571. This work was initiated at a Cell Biology Hackathon supported by NSF grant MCB 1411898 to Wallace Marshall.

No conflicts of interest declared.

## REFERENCES

1. Liu, S., Jobert, F., Rahneshan, Z., Doyle, S.M., and Robert, S. (2021). Solving the Puzzle of Shape Regulation in Plant Epidermal Pavement Cells. Annu Rev Plant Biol 72, 525–550.

2. Eng, R.C., and Sampathkumar, A. (2018). Getting into shape: the mechanics behind plant morphogenesis. Curr Opin Plant Biol 46, 25–31.

3. Trinh, D.-C., Alonso-Serra, J., Asaoka, M., Colin, L., Cortes, M., Malivert, A., Takatani, S., Zhao, F., Traas, J., Trehin, C., et al. (2021). How Mechanical Forces Shape Plant Organs. Curr Biol 31, R143–R159.

4. Smithers, E.T., Luo, J., and Dyson, R.J. (2019). Mathematical principles and models of plant growth mechanics: from cell wall dynamics to tissue morphogenesis. J Exp Bot 70, 3587–3600.

5. Long, Y., Cheddadi, I., Mosca, G., Mirabet, V., Dumond, M., Kiss, A., Traas, J., Godin, C., and Boudaoud, A. (2020). Cellular Heterogeneity in Pressure and Growth Emerges from Tissue Topology and Geometry. Curr Biol 30, 1504–1516.e8.

6. Ali, O., Oliveri, H., Traas, J., and Godin, C. (2019). Simulating Turgor-Induced Stress Patterns in Multilayered Plant Tissues. Bull Math Biol 81, 3362–3384.

7. Pacini, E., Guarnieri, M., and Nepi, M. (2006). Pollen carbohydrates and water content during development, presentation, and dispersal: a short review. Protoplasma 228, 73–77.

8. Rozier, F., Riglet, L., Kodera, C., Bayle, V., Durand, E., Schnabel, J., Gaude, T., and Fobis-Loisy, I. (2020). Live-cell imaging of early events following pollen perception in self-incompatible Arabidopsis thaliana. J Exp Bot 71, 2513–2526.

9. Zheng, Y.-Y., Lin, X.-J., Liang, H.-M., Wang, F.-F., and Chen, L.-Y. (2018). The Long Journey of Pollen Tube in the Pistil. International Journal of Molecular Sciences 19, 3529.

10. Lohani, N., Singh, M.B., and Bhalla, P.L. (2020). High temperature susceptibility of sexual reproduction in crop plants. J Exp Bot 71, 555–568.

11. Hill, A.E., Shachar-Hill, B., Skepper, J.N., Powell, J., and Shachar-Hill, Y. (2012). An osmotic model of the growing pollen tube. PLoS One 7, e36585.

12. Liu, J., and Hussey, P.J. (2014). Dissecting the regulation of pollen tube growth by modeling the interplay of hydrodynamics, cell wall and ion dynamics. Front Plant Sci 5.

13. Hemelryck, M.V., Bernal, R., Ispolatov, Y., and Dumais, J. (2018). Lily Pollen Tubes Pulse According to a Simple Spatial Oscillator. Scientific Reports 8.

14. Tian, W., Wang, C., Gao, Q., Li, L., and Luan, S. (2020). Calcium spikes, waves and oscillations in plant development and biotic interactions. Nat Plants 6, 750–759.

15. Vogler, H., Draeger, C., Weber, A., Felekis, D., Eichenberger, C., Routier-Kierzkowska, A.-L., Boisson-Dernier, A., Ringli, C., Nelson, B.J., Smith, R.S., et al. (2013). The pollen tube: a soft shell with a hard core. Plant J 73, 617–627.

16. Fayant, P., Girlanda, O., Chebli, Y., Aubin, C.-É., Villemure, I., and Geitmann, A. (2010). Finite Element Model of Polar Growth in Pollen Tubes. The Plant Cell 22, 2579–2593.

17. Katifori, E., Alben, S., Cerda, E., Nelson, D.R., and Dumais, J. (2010). Foldable structures and the natural design of pollen grains. Proc Natl Acad Sci U S A 107, 7635–7639.

18. Božič, A., and Šiber, A. (2020). Mechanical design of apertures and the infolding of pollen grain. Proc Natl Acad Sci U S A 117, 26600–26607.

19. Mayfield, J.A., and Preuss, D. (2000). Rapid initiation of Arabidopsis pollination requires the oleosin-domain protein GRP17. Nat Cell Biol 2, 128–130.

20. Updegraff, E.P., Zhao, F., and Preuss, D. (2009). The extracellular lipase EXL4 is required for efficient hydration of Arabidopsis pollen. Sex Plant Reprod 22, 197–204.

21. Wang, L., Clarke, L.A., Eason, R.J., Parker, C.C., Qi, B., Scott, R.J., and Doughty, J. (2017). PCP-B class pollen coat proteins are key regulators of the hydration checkpoint in Arabidopsis thaliana pollen-stigma interactions. New Phyt 213, 764–777.

22. Hülskamp, M., Kopczak, S.D., Horejsi, T.F., Kihl, B.K., and Pruitt, R.E. (1995). Identification of genes required for pollen-stigma recognition in Arabidopsis thaliana. Plant J 8, 703–714.

23. Gao, X.-Q., Liu, C.Z., Li, D.D., Zhao, T.T., Li, F., Jia, X.N., Zhao, X.-Y., and Zhang, X.S. (2016). The Arabidopsis KINβγ Subunit of the SnRK1 Complex Regulates Pollen Hydration on the Stigma by Mediating the Level of Reactive Oxygen Species in Pollen. PLoS Genet 12, e1006228.

24. Li, D.-D., Guan, H., Li, F., Liu, C.-Z., Dong, Y.-X., Zhang, X.-S., and Gao, X.-Q. (2017). Arabidopsis shaker pollen inward K+ channel SPIK functions in SnRK1 complex-regulated pollen hydration on the stigma. J Integr Plant Biol 59, 604–611.

25. Moon, S., and Jung, K.-H. (2020). First Steps in the Successful Fertilization of Rice and Arabidopsis: Pollen Longevity, Adhesion and Hydration. Plants (Basel) 9, E956.

26. Edlund, A.F. (2004). Pollen and Stigma Structure and Function: The Role of Diversity in Pollination. Plant Cell 16, S84–S97.

27. Wang, R., and Dobritsa, A.A. (2018). Exine and Aperture Patterns on the Pollen Surface: Their Formation and Roles in Plant Reproduction. Annual Plant Rev Online. 1, 1–40.

28. Vieira, A.M., and Feijó, J.A. (2016). Hydrogel control of water uptake by pectins during in vitro pollen hydration of Eucalyptus globulus. Am J Bot 103, 437–451.

29. Fan, T.-F., Park, S., Shi, Q., Zhang, X., Liu, Q., Song, Y., Chin, H., Ibrahim, M.S.B., Mokrzecka, N., Yang, Y., et al. (2020). Transformation of hard pollen into soft matter. Nat Comm 11.

30. Leroux, C., Bouton, S., Kiefer-Meyer, M.-C., Fabrice, T.N., Mareck, A., Guénin, S., Fournet, F., Ringli, C., Pelloux, J., Driouich, A., et al. (2015). PECTIN METHYLESTERASE48 Is Involved in Arabidopsis Pollen Grain Germination. Plant Phys 167, 367–380.

31. Windari, E.A., Ando, M., Mizoguchi, Y., Shimada, H., Ohira, K., Kagaya, Y., Higashiyama, T., Takayama, S., Watanabe, M., and Suwabe, K. (2021). Two aquaporins, SIP1;1 and PIP1;2, mediate water transport for pollen hydration in the Arabidopsis pistil. Plant Biotechnol (Tokyo) 38, 77–87.

32. Hamilton, E.S., Jensen, G.S., Maksaev, G., Katims, A., Sherp, A.M., and Haswell, E.S. (2015). Mechanosensitive channel MSL8 regulates osmotic forces during pollen hydration and germination. Science 350, 438–441.

33. Poole, K. (2021). The Diverse Physiological Functions of Mechanically Activated Ion Channels in Mammals. Annu Rev Physiol.

34. Cox, C.D., Bavi, N., and Martinac, B. (2018). Bacterial Mechanosensors. Annu Rev Physiol 80, 71–93.

35. Codjoe, J.M., Miller, K., and Haswell, E.S. (2021). Plant Cell Mechanobiology: Greater Than the Sum of its Parts. Plant Cell. in press.

36. Martinac, B., Buechner, M., Delcour, A.H., Adler, J., and Kung, C. (1987). Pressure-sensitive ion channel in Escherichia coli. Proc Natl Acad Sci U S A 84, 2297–2301.

37. Sukharev, S.I., Blount, P., Martinac, B., Blattner, F.R., and Kung, C. (1994). A large-conductance mechanosensitive channel in E. coli encoded by mscL alone. Nature 368, 265–268.

38. Sukharev, S. (2002). Purification of the small mechanosensitive channel of Escherichia coli (MscS): the subunit structure, conduction, and gating characteristics in liposomes. Biophys J 83, 290–298.

39. Berrier, C., Coulombe, A., Szabo, I., Zoratti, M., and Ghazi, A. (1992). Gadolinium ion inhibits loss of metabolites induced by osmotic shock and large stretch-activated channels in bacteria. Eur J Biochem 206, 559–565.

40. Levina, N., Tötemeyer, S., Stokes, N.R., Louis, P., Jones, M.A., and Booth, I.R. (1999). Protection of Escherichia coli cells against extreme turgor by activation of MscS and MscL mechanosensitive channels: identification of genes required for MscS activity. EMBO J 18, 1730–1737.

41. Buda, R., Liu, Y., Yang, J., Hegde, S., Stevenson, K., Bai, F., and Pilizota, T. (2016). Dynamics of Escherichia coli’s passive response to a sudden decrease in external osmolarity. Proc Natl Acad Sci U S A 113, E5838–E5846.

42. Hamilton, E.S., and Haswell, E.S. (2017). The Tension-sensitive Ion Transport Activity of MSL8 is Critical for its Function in Pollen Hydration and Germination. Plant & Cell Physiol 58, 1222–1237.

43. Basu, D., and Haswell, E.S. (2017). Plant mechanosensitive ion channels: an ocean of possibilities. Curr Opin Plant Biol 40, 43–48.

44. Veley, K.M., Maksaev, G., Frick, E.M., January, E., Kloepper, S.C., and Haswell, E.S. (2014). Arabidopsis MSL10 has a regulated cell death signaling activity that is separable from its mechanosensitive ion channel activity. Plant Cell 26, 3115–3131.

45. Maksaev, G., Shoots, J.M., Ohri, S., and Haswell, E.S. (2018). Nonpolar residues in the presumptive pore-lining helix of mechanosensitive channel MSL10 influence channel behavior and establish a nonconducting function. Plant Direct 2, e00059.

46. Wang, Y., Coomey, J., Miller, K., Jensen, G.S., and Haswell, E.S. (2021). Interactions between a mechanosensitive channel and cell wall integrity signaling influence pollen germination in Arabidopsis thaliana. https://www.biorxiv.org/content/10.1101/2021.08.24.457556v1

47. Nezhad, A.S., Naghavi, M., Packirisamy, M., Bhat, R., and Geitmann, A. (2013). Quantification of the Young’s modulus of the primary plant cell wall using Bending-Lab-On-Chip (BLOC). Lab Chip 13, 2599–2608.

48. Wolfe, J., and Steponkus, P.L. (1983). Mechanical Properties of the Plasma Membrane of Isolated Plant Protoplasts: Mechanism of Hyperosmotic and Extracellular Freezing Injury. Plant Physiol. 71, 276–285.

49. Pertl, H., Pockl, M., Blaschke, C., and Obermeyer, G. (2010). Osmoregulation in Lilium Pollen Grains Occurs via Modulation of the Plasma Membrane H+ ATPase Activity by 14-3-3 Proteins. Plant Physiol. 154, 1921–1928.

50. Le Roux, A.-L., Quiroga, X., Walani, N., Arroyo, M., and Roca-Cusachs, P. (2019). The plasma membrane as a mechanochemical transducer. Phil Trans Royal Soc B: Biol Sci 374, 20180221.

51. Talemi, S.R., Tiger, C.-F., Andersson, M., Babazadeh, R., Welkenhuysen, N., Klipp, E., Hohmann, S., and Schaber, J. (2016). Systems Level Analysis of the Yeast Osmo-Stat. Sci Rep 6, 30950.

52. Altenburg, T., Goldenbogen, B., Uhlendorf, J., and Klipp, E. (2019). Osmolyte homeostasis controls single-cell growth rate and maximum cell size of Saccharomyces cerevisiae. NPJ Syst Biol Appl 5, 34.

53. Platt-Aloia, K.A., Lord, E.M., DeMason, D.A., and Thomson, W.W. (1986). Freeze-fracture observations on membranes of dry and hydrated pollen from Collomia, Phoenix and Zea. Planta 168, 291–298.

54. Fruleux, A., Verger, S., and Boudaoud, A. (2019). Feeling Stressed or Strained? A Biophysical Model for Cell Wall Mechanosensing in Plants. Front Plant Sci 10.

55. Boer, M., Anishkin, A., and Sukharev, S. (2011). Adaptive MscS gating in the osmotic permeability response in E. coli: the question of time. Biochem 50, 4087–4096.

56. Kamaraju, K., Belyy, V., Rowe, I., Anishkin, A., and Sukharev, S. (2011). The pathway and spatial scale for MscS inactivation. J Gen Physiol 138, 49–57.

57. Paraschiv, A., Hegde, S., Ganti, R., Pilizota, T., and Šarić, A. (2020). Dynamic Clustering Regulates Activity of Mechanosensitive Membrane Channels. Phys Rev Lett 124, 048102.

58. Tran, D., Girault, T., Guichard, M., Thomine, S., Leblanc-Fournier, N., Moulia, B., de Langre, E., Allain, J.-M., and Frachisse, J.-M. (2021). Cellular transduction of mechanical oscillations in plants by the plasma-membrane mechanosensitive channel MSL10. Proc Natl Acad Sci U S A 118, e1919402118.

59. Maksaev, G., and Haswell, E.S. (2012). MscS-Like10 is a stretch-activated ion channel from Arabidopsis thaliana with a preference for anions. Proc Natl Acad Sci U S A 109, 19015–19020.

60. Schiller, L.R., Emmett, M., Santa Ana, C.A., and Fordtran, J.S. (1988). Osmotic effects of polyethylene glycol. Gastroenterology 94, 933–941.

61. Twell, D., Yamaguchi, J., and McCormick, S. (1990). Pollen-specific gene expression in transgenic plants: coordinate regulation of two different tomato gene promoters during microsporogenesis. Development 109, 705–713.

62. Elleman, C.J., and Dickinson, H.G. (1986). Pollen-stigma interactions in Brassica. IV. Structural reorganization in the pollen grains during hydration. J Cell Sci 80, 141–157.

63. Heslop-Harrison, J. (1979). Pollen Walls as Adaptive Systems. Annals of the Missouri Botanical Garden 66, 813.

64. Shivanna, K.R., and Heslop-Harrison, J. (1981). Membrane State and Pollen Viability. Annals of Botany 47, 759–770.

65. Kerhoas, C., Gay, G., and Dumas, C. (1987). A multidisciplinary approach to the study of the plasma membrane of Zea mays pollen during controlled dehydration. Planta 171, 1–10.

66. Groulx, N., Boudreault, F., Orlov, S.N., and Grygorczyk, R. (2006). Membrane reserves and hypotonic cell swelling. J Membr Biol 214, 43–56.

67. Zhang, Y., and Hamill, O.P. (2000). On the discrepancy between whole-cell and membrane patch mechanosensitivity in Xenopus oocytes. J Physiol 523 Pt 1, 101–115.

68. Gauthier, N.C., Fardin, M.A., Roca-Cusachs, P., and Sheetz, M.P. (2011). Temporary increase in plasma membrane tension coordinates the activation of exocytosis and contraction during cell spreading. Proc Natl Acad Sci U S A 108, 14467–14472.

69. Thompson, D.S., and Islam, A. (2021). Plant Cell Wall Hydration and Plant Physiology: An Exploration of the Consequences of Direct Effects of Water Deficit on the Plant Cell Wall. Plants (Basel) 10, 1263.

70. Lee, C.P., Maksaev, G., Jensen, G.S., Murcha, M.W., Wilson, M.E., Fricker, M., Hell, R., Haswell, E.S., Millar, A.H., and Sweetlove, L.J. (2016). MSL1 is a mechanosensitive ion channel that dissipates mitochondrial membrane potential and maintains redox homeostasis in mitochondria during abiotic stress. Plant J 88, 809–825.

71. Anderson, C.T., and Kieber, J.J. (2020). Dynamic Construction, Perception, and Remodeling of Plant Cell Walls. Ann Rev Plant Biol 71, 39–69.

72. Cannon, M.C., Terneus, K., Hall, Q., Tan, L., Wang, Y., Wegenhart, B.L., Chen, L., Lamport, D.T.A., Chen, Y., and Kieliszewski, M.J. (2008). Self-assembly of the plant cell wall requires an extensin scaffold. Proc Natl Acad Sci U S A 105, 2226–2231.

73. Schindelin, J., Arganda-Carreras, I., Frise, E., Kaynig, V., Longair, M., Pietzsch, T., Preibisch, S., Rueden, C., Saalfeld, S., Schmid, B., et al. (2012). Fiji: an open-source platform for biological-image analysis. Nat Methods 9, 676–682.

