## Supplementary Files for "In vitro experiments and kinetic models of pollen hydration show that MSL8 is not a simple tension-gated osmoregulator"

### SUPPLEMENTAL FIGURES

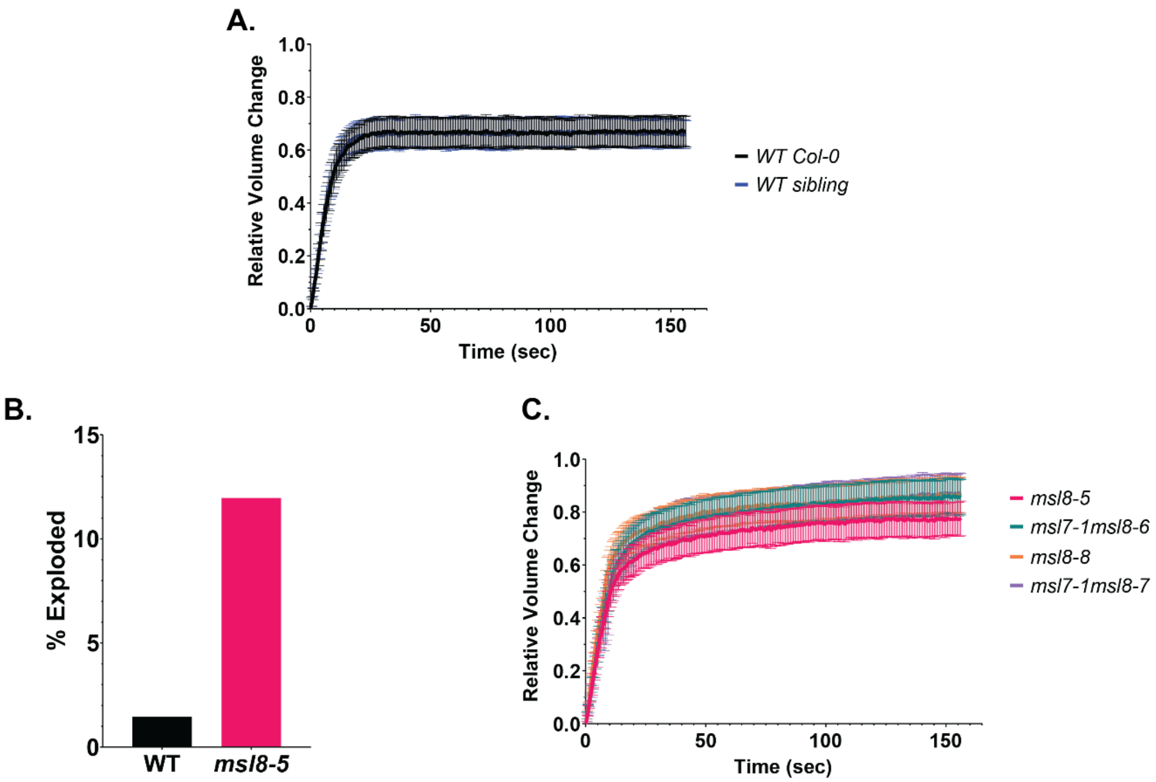


**Supplemental Figure 1: In vitro pollen hydration curves of all *msl8* mutant and WT lines.**
(A) Hydration curves for two wild type lines. WT Col-0 is the original parent line and WT sibling was segregated from the *msl7-1msl8-7* line. N = 30 grains for each genotype; Bars are 95% confidence intervals (CI). (B) Percentage of pollen grains that visibly burst during the first 150 seconds of hydration (N = 206 grains for WT; N = 234 grains for *msl8-5*). (C) Hydration curves from all *msl8* mutant lines.

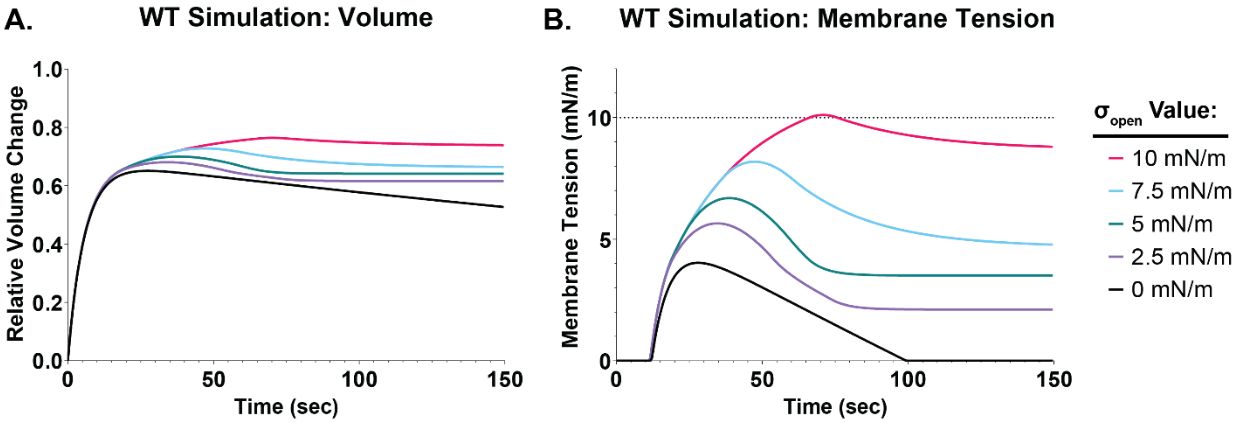


**Supplemental Figure 2: Effects of altering the channel opening membrane tension threshold.** (A) Volume output of simulations testing $\sigma_{open}$ values of 0, 2.5, 5, 7.5, and 10 mN/m. Note that 5 mN/m is the estimated value based on existing literature. (B) Membrane tension output from the same five simulations. The dashed line marks the estimated lytic tension of the membrane.

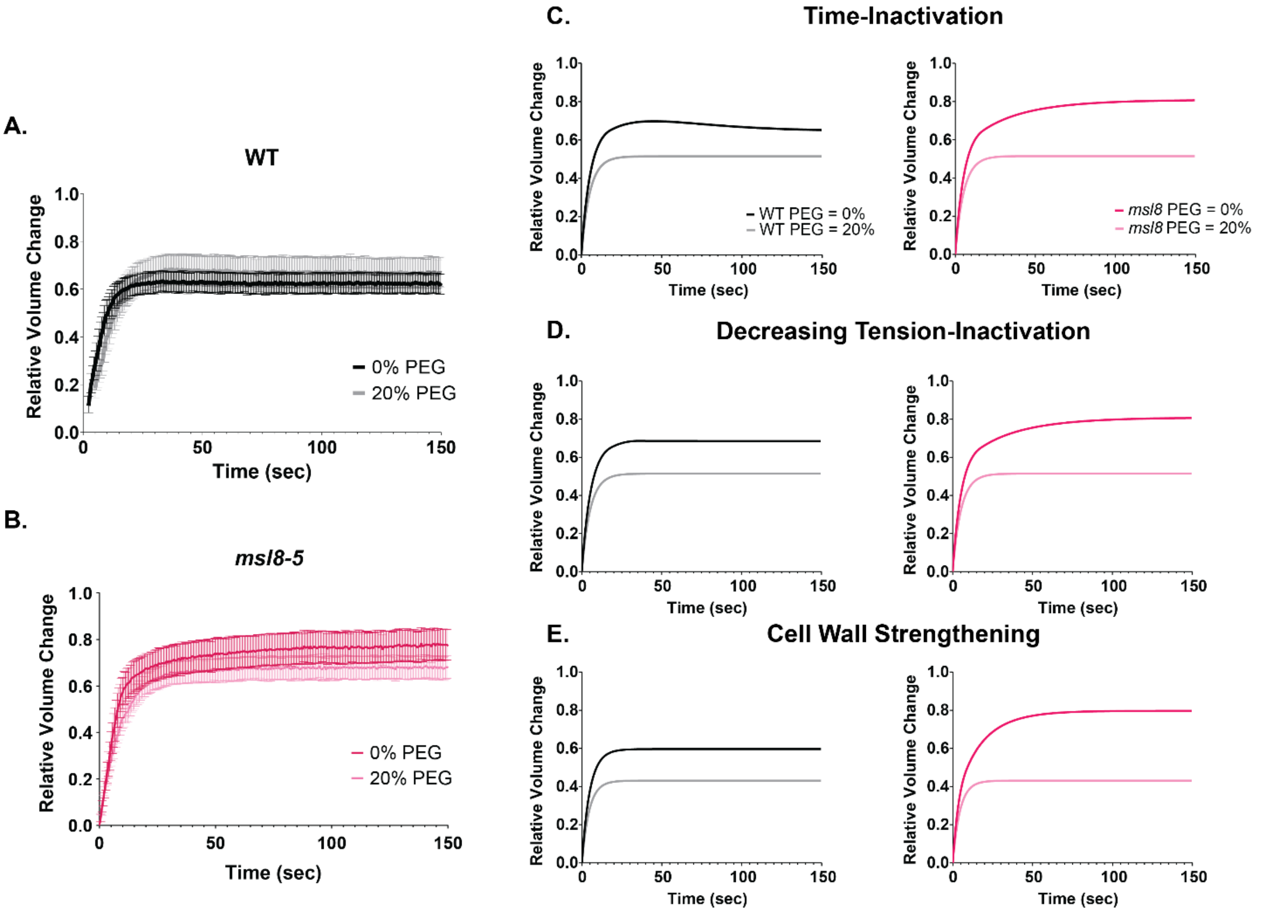


**Supplemental Figure 3: Hydration in an osmolyte-rich solution experimental observations and simulations.**
Relative volume change of WT (A) and *msl8-5* (B) pollen hydrated in water or 20% PEG 3350 solution. N = 30 pollen grains for each genotype/treatment. Bars are 95% CI. Simulation results for both WT (left) and *msl8* (right) hydration in 0% PEG and 20% PEG (assuming the MSL8 channels inactivate after a period time (C), MSL8 channels inactivate when tension starts to decrease (D), or the presence of the MSL8 channels strengthens the cell wall (E).

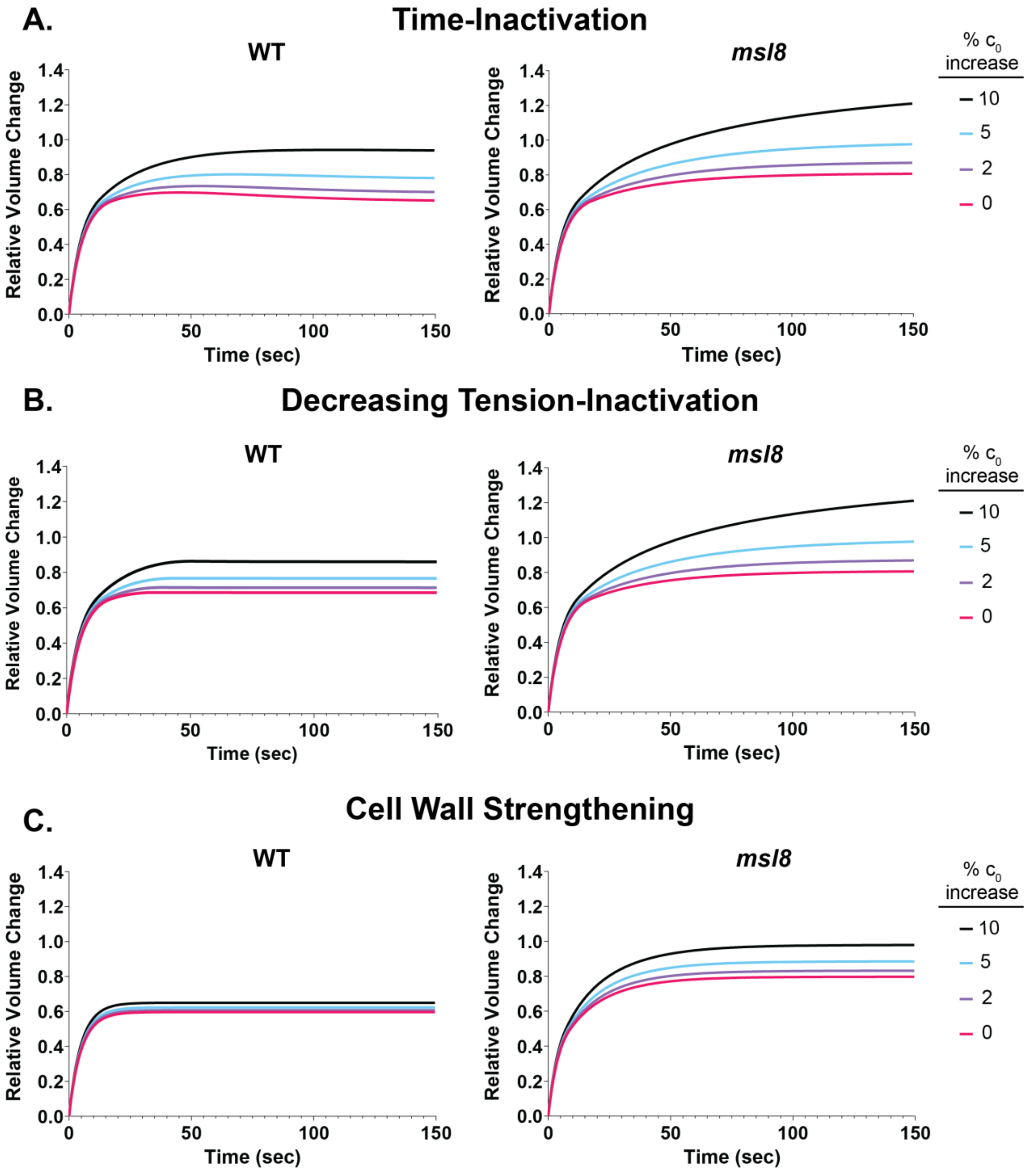


**Supplemental Figure 4: Hydration simulations of pollen with increased cytoplasmic osmolarity.**

Simulation results for both WT (left) and *msl8* (right) pollen grains with increasing initial osmotic concentration values assuming either the MSL8 channels inactivate after a period time (A), MSL8 channels inactivate when tension starts to decrease (B), or the presence of the MSL8 channels strengthens the cell wall (C).

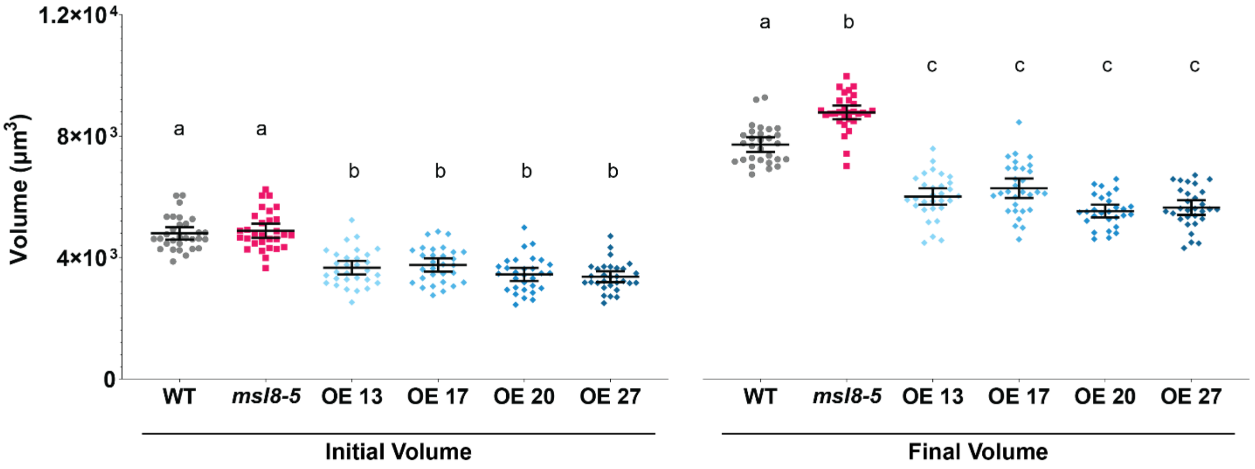


**Supplemental Figure 5: Volume of pollen overexpressing MSL8-YFP.**
Initial pollen volume (left) and final volume (right) of WT, *msl8-5*, and four MSL8-YFP overexpression lines (OE) (N = 30 grains for each). Bars are mean with 95% CI. One-way ANOVA test with multiple comparisons was used to compare each genotype among initial volume and final volume groups. Letters indicate a significant difference ($p < 0.05$). Grubbs test for outliers was conducted on all genotypes. One outlier from WT final volume and two outliers from OE 20 final volume were removed. Excluding the outliers did not affect the ANOVA results.

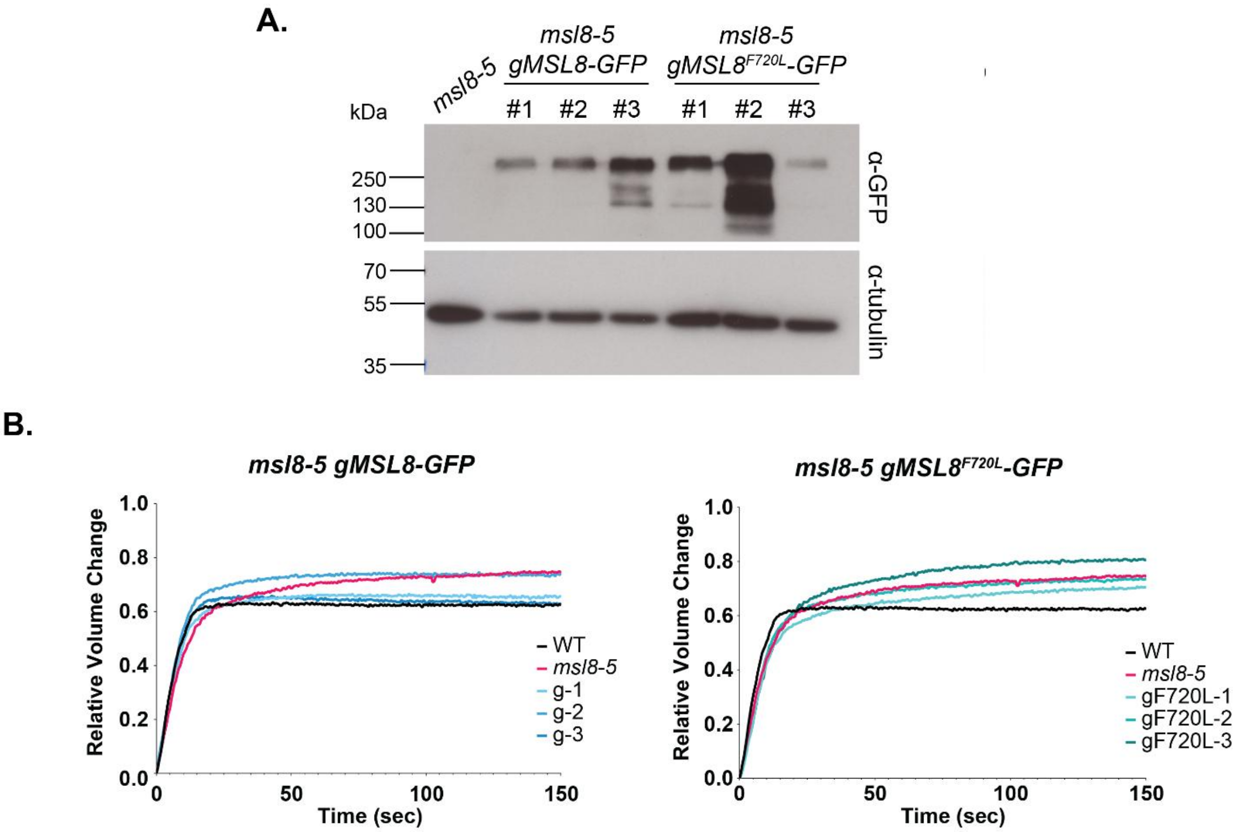


**Supplemental Figure 6: Characterization of *MSL8* lines.**
(A) Immunoblot confirmation of protein presence in *msl8-5* lines expressing genomic MSL8-GFP or MSL8-GFP with the point mutation, F720L, which eliminates channel function. Protein blot was probed with anti-GFP (top). The blot was then stripped and re-probed with anti-tubulin (bottom). (B) Complete hydration curves of the lines.

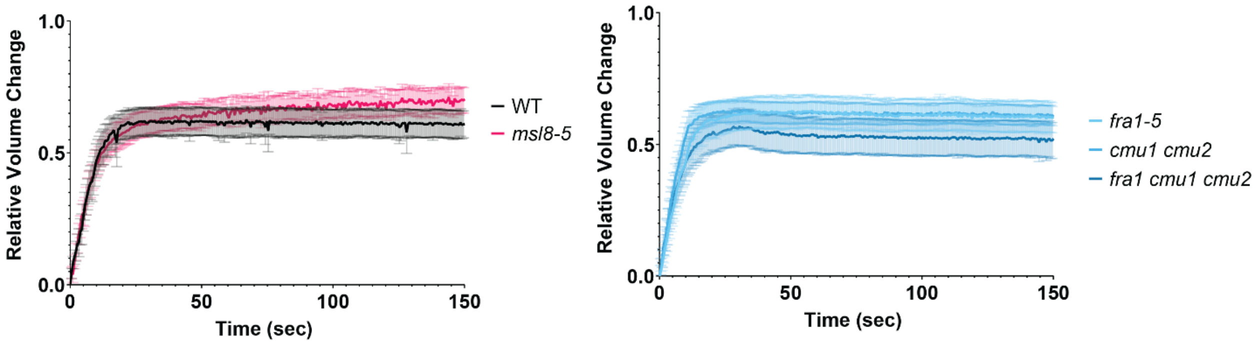


**Supplemental Figure 7: Hydration of pollen cell wall mutants.**
Hydration curves of WT and *msl8-5* controls (left) and three cell wall mutants (*fra1-5*, *cmu1 cmu2*, and *fra1-5 cmu1 cmu2*) (right) (N=30 grains per genotype). Bars are 95% CI. All hydrations were performed in 10% PEG due to the high rate of explosions of *fra1 cmu1 cmu2* pollen.
